# Mechanical strain stimulates COPII-dependent trafficking via Rac1

**DOI:** 10.1101/2022.01.23.477215

**Authors:** Santosh Phuyal, Elena Djaerff, Anabel-Lise Le Roux, Martin J. Baker, Daniela Fankhauser, Sayyed Jalil Mahdizadeh, Veronika Reiterer, Jennifer C. Kahlhofer, David Teis, Marcelo G. Kazanietz, Stephan Geley, Leif Eriksson, Pere Roca-Cusachs, Hesso Farhan

## Abstract

Secretory trafficking from the endoplasmic reticulum (ER) is subject to regulation by extrinsic and intrinsic factors. While much of the focus has been on biochemical triggers, little is known whether and how the ER is subject to regulation by mechanical signals. Here, we show that COPII-dependent ER-export is regulated by mechanical strain. Mechanotransduction to the ER was mediated via a previously unappreciated ER-localized pool of the small GTPase Rac1. Mechanistically, we show that Rac1 interacts with the small GTPase Sar1 to drive budding of COPII carriers and stimulate ER-to-Golgi transport. Altogether, we establish an unprecedented link between mechanical strain and export from the ER.

## Introduction

Endoplasmic reticulum (ER) exit sites (ERES) are specialized ribosome-free domains of the rough ER (Farhan *et al*, 2007; Malkus *et al*, 2002; Shomron *et al*, 2021), which give rise to an intricate network of tubules and vesicles that ferry cargo towards distal compartments (Phuyal & Farhan, 2021; Weigel *et al*, 2021; Zeuschner *et al*, 2006). In recent years, ERES have emerged as platforms that integrate signaling pathways in response to alterations of secretory protein load, starvation, or mitogens (Centonze & Farhan, 2019; Centonze *et al*, 2019b; Farhan *et al*, 2008; Farhan *et al*, 2010; Subramanian *et al*, 2019; Zacharogianni *et al*, 2011). Thus, signaling to ERES allows cells to tune the secretory rate to meet the changing requirements during cell growth and proliferation. At ERES, secretory proteins leave the ER via COPII-dependent carriers. The assembly of the COPII coat is initiated by the small GTPase Sar1, which is activated by its exchange factor Sec12, a transmembrane ER-resident protein. Recruitment of Sec12 was shown to reconstitute a critical event in the biogenesis of ERES (Maeda *et al*, 2017). Moreover, we showed recently that phosphorylation of Sec12 regulates ERES number (Centonze *et al*, 2019a).

Besides intracellular and environmental chemical stimuli, cells are constantly exposed to mechanical stimuli such as substrate stiffness, compression, or tensile forces (Discher *et al*, 2005). It is now well established that such mechanical cues trigger signaling pathways that mediate changes in cell differentiation, proliferation, growth and survival (Engler *et al*, 2006; Gudipaty *et al*, 2017; Janmey *et al*, 2020; Roca-Cusachs *et al*, 2013). Most research in this area has focused on the plasma membrane, the nucleus or the cytoskeleton as receivers and mediators of mechanical signaling (Phuyal & Baschieri, 2020). However, it is currently poorly understood whether and how mechanical stress has any effect on secretory pathway compartments such as ERES.

The small GTPase Rac1 is a major regulator of actin cytoskeleton remodeling (Nobes & Hall, 1995). Rac1 has been reported to orchestrate spatially restricted signaling cascades at various subcellular organelles (Payapilly & Malliri, 2018; Phuyal & Farhan, 2019). However, any presence of functionally active Rac1 at the ER remains unexplored. Importantly, previous reports show that Rac1 signals downstream of mechanical cues to regulate cellular proliferation, gene expression, nutrient transport, and epithelial wound healing (Desai *et al*, 2008; Gould *et al*, 2016; Katsumi *et al*, 2002; Kumar *et al*, 2004; Liu *et al*, 2007; Verma *et al*, 2011; Yamane *et al*, 2007). An open question is whether Rac1 signaling regulates the early secretory pathway, and whether mechanical cues act as a stimulus for this process.

In this study, we demonstrate that the early secretory pathway responds to mechanical cues. Our results show that changes in mechanical tension increases ERES number and enhances the rate of ER-exit in a manner dependent on Rac1. We further show that Rac1 regulates ERES by interacting with the small GTPase Sar1.

## Results

### Mechanical strain stimulates the early secretory pathway in a Rac1-dependent manner

We initially opted for growing HeLa cells on fibronectin coated micropatterned surface that contained multiple geometric shapes of two different sizes (small: 700 µm^2^, large: 1600 µm^2^) to impose mechanical tension on cells (Albert & Schwarz, 2014). Cells were allowed to accommodate to the patterns of different sizes for four hours and were fixed, labelled with anti-Sec31 as an ERES marker and analyzed by fluorescence microscopy. Strikingly, forcing cells to occupy a larger surface area led to an increased ERES number irrespective of the geometry (**Fig.1Aleft panel, B, Fig.S1A-B**). It appears intuitive that larger cells have more ERES, because they likely have a larger proteome and thus a larger secretome. However, this does not apply to cells grown on larger patterns, which are forced to occupy a larger surface area for a short duration (4 hours) and are unlikely to have larger volumes and a larger biomass. Thus, the increase ERES is not a consequence of larger biomass, which opened the possibility that ERES respond to mechanical tension. Next, we aimed to demonstrate the effect of mechanical tension in live cells. To this end, we cultured HeLa cells transiently expressing GFP-tagged Sec16A on a fibronectin coated poly(dimethylsiloxane) (PDMS) membrane and subjected them to 7.5% biaxial stretch. We noted that stretching cells resulted in a rapid increase in the number of ERES (**Fig.1C-D, movieS1 upper panel)**. We also examined the response of ERES when HeLa cells were exposed to acute (5 min) mechanical strain through hypotonic swelling and observed an increase in ERES (**Fig. S1C-D**). Thus, mechanical tension results in a marked upregulation in the number of ERES.

**Figure 1.**
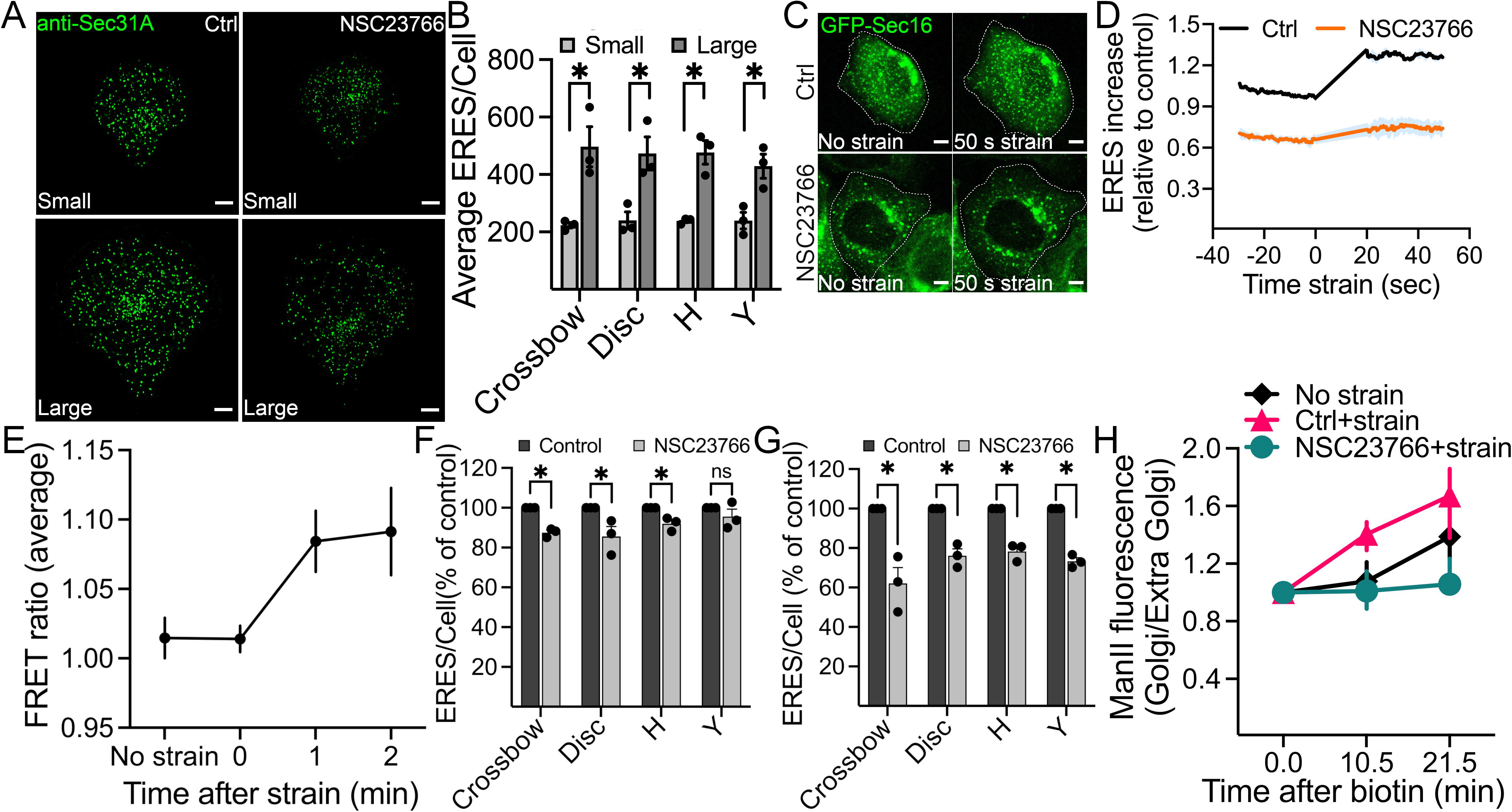
Mechanical strain stimulates ERES and ER-to-Golgi transport. (A) Representative immunofluorescence images of Sec31A marked ERES in HeLa cells cultured on crossbow shaped micropatterned surface of small (upper panel) or large (lower panel) size for 4 hours. Cells were treated with DMSO (Ctrl) or 50 µM NSC23766 (Rac1 inhibitor) for 4 hours prior to fixation and immunostaining. Scale bars: 5 µm (B) The number of ERES per cell for each pattern size was quantified from at least 25 cells/per condition from 3 independent experiments. Error bars represent SEM. (C) HeLa cells transiently expressing GFP-Sec16A were grown on PDMS membranes, treated as indicated and subjected to 7.5% equibiaxial stretch. The effect of stretch on ERES was then monitored by live cell imaging. Representative still images are shown. Scale bar: 5 µm. See also movieS1. (D) Quantification of the experiments in (C). ERES increase in HeLa cells before and during equibiaxial stretch were quantified from a total of 19 cells in three live cell imaging experiments. Baseline GFP-Sec16 count from control (Ctrl) before strain was used to normalize GFP-Sec16 counts for each condition. Shading represents SEM. (E) HeLa cells transfected with Rac1 FRET biosensor were cultured on PDMS membranes, subjected to 7.5% equibiaxial stretch and Rac1 FRET ratio was measured for indicated time points. Graph shows average FRET ratio calculated from a total from 16 cells from 3 independent experiments. (**F-G**) Quantification of ERES in HeLa cells seeded on small (F) and large (G) micropatterned surfaces (described in A). Data is derived from 90-100 cells from three independent experiments. Error bars show SEM. (**H**) Golgi arrival kinetics of GFP-ManII-RUSH in HeLa cells stably expressing Str-KDEL- SBP-GFP-ManII (RUSH system). Cell were grown on PDMS membranes, treated as indicated and subjected to 7.5% equibiaxial stretch. See also movieS2. ManII arrival at the Golgi was monitored in nineteen cells in two independent experiments. Error bars indicate standard deviation. See also movieS2. In all graphs, asterisks (*) mark statistical significance (p-value < 0.05; Student’s unpaired *t* test).

We next asked how mechanical strain is linked to the early secretory pathway. Rac1 activation and signaling has been previously linked to mechanical strain (Katsumi *et al*., 2002). To validate that Rac1 is activated in our experimental system upon cell stretch, we used cells expressing a Rac1 FRET biosensor (Fritz *et al*, 2013). We observed a clear increase in Rac1 activation upon equibiaxial stretching (**Fig.1E**). To test whether Rac1 is involved in the stretch-induced increase in ERES, we cultured HeLa cells on the micropatterned surface, allowed them to attach for 30 min and then treated with the Rac1 inhibitor NSC23766 for four hours. NSC23766 is a reversible Rac1-specific inhibitor that occupies the GEF-recognition groove centering on Trp56 of Rac1 to inhibit its activation by some GEFs, such as TIAM1 (Gao *et al*, 2004). Rac1 inhibition resulted in a decrease in the number of ERES. Notably, the reduction of ERES number was more pronounced in cells that occupied larger geometries compared to the cells occupying smaller geometries (**Fig.1Aright panel, F-G**). In a similar manner, NSC23766 also blocked the acute increase in ERES number upon biaxial stretching (**Fig.1C-D, movieS1**). These effects were not due to an effect of NSC23766 on cell surface area (**Fig.S1E-G**). Thus, the increase in ERES number by mechanical stretch is dependent on Rac1.

Because ERES are sites for cargo exit from the ER, we tested whether mechanically challenged cells would exhibit a change in the rate of ER-to-Golgi transport. Therefore, we exploited the retention using selective hook (RUSH) assay (Boncompain *et al*, 2012) with Mannosidase II (ManII) as secretory reporter. We applied biaxial strain to HeLa cells stably expressing the RUSH reporter ManII and observed an enhanced ER-export of ManII in stretched cells compared to that in non-stretched cells. Again, Rac1 was essential for stretch-induced acceleration of ER-export since perturbation of its activity with the inhibitor NSC23766 delayed the ER-to-Golgi transport (**Fig.1H, movieS2**).

Based on the findings from these experiments, we conclude that the early secretory pathway responds to mechanical strain and this response depends on Rac1 signaling activity.

### The early secretory pathway is required for cellular adaptation to mechanical strain

Having established the first link between mechanical strain and the early secretory pathway, we examined whether the early secretory pathway has any functional consequences for cellular adaptation during mechanical stimulation. For this purpose, we disrupted ERES assembly in cells using small interfering RNA (siRNA) targeting the small GTPases Sar1A and Sar1B. We verified depletion of Sar1 level using Western blot (Fig.S2A). Silencing Sar1A/B expression disrupts the assembly and the functionality of ERES(Cutrona *et al*, 2013). We cultured Sar1-depleted HeLa cells on the micropatterned surface, allowed them to adapt for 3 hours, and monitored whether cells occupy the whole geometry. We found that a significantly large proportion of Sar1-depleted cells failed to fully spread to cover the large micropattern geometries compared to the control cells (Fig. 2A-B). As expected, for cells covering the smaller micropattern geometries, we did not observe any difference (Fig. S2B-C). This observation suggests that the dysfunctional ERES impacts cellular ability to adapt to mechanical stimulation. Of note, the defect in cellular adaptation was not due to an overall decrease in cell size since depletion of Sec16A, another key protein required for ERES function, did not reduce cell size (Fig. 2C). These findings support the idea that the early secretory pathway plays an important role during cellular adaptation to mechanical tension.

**Figure 2.**
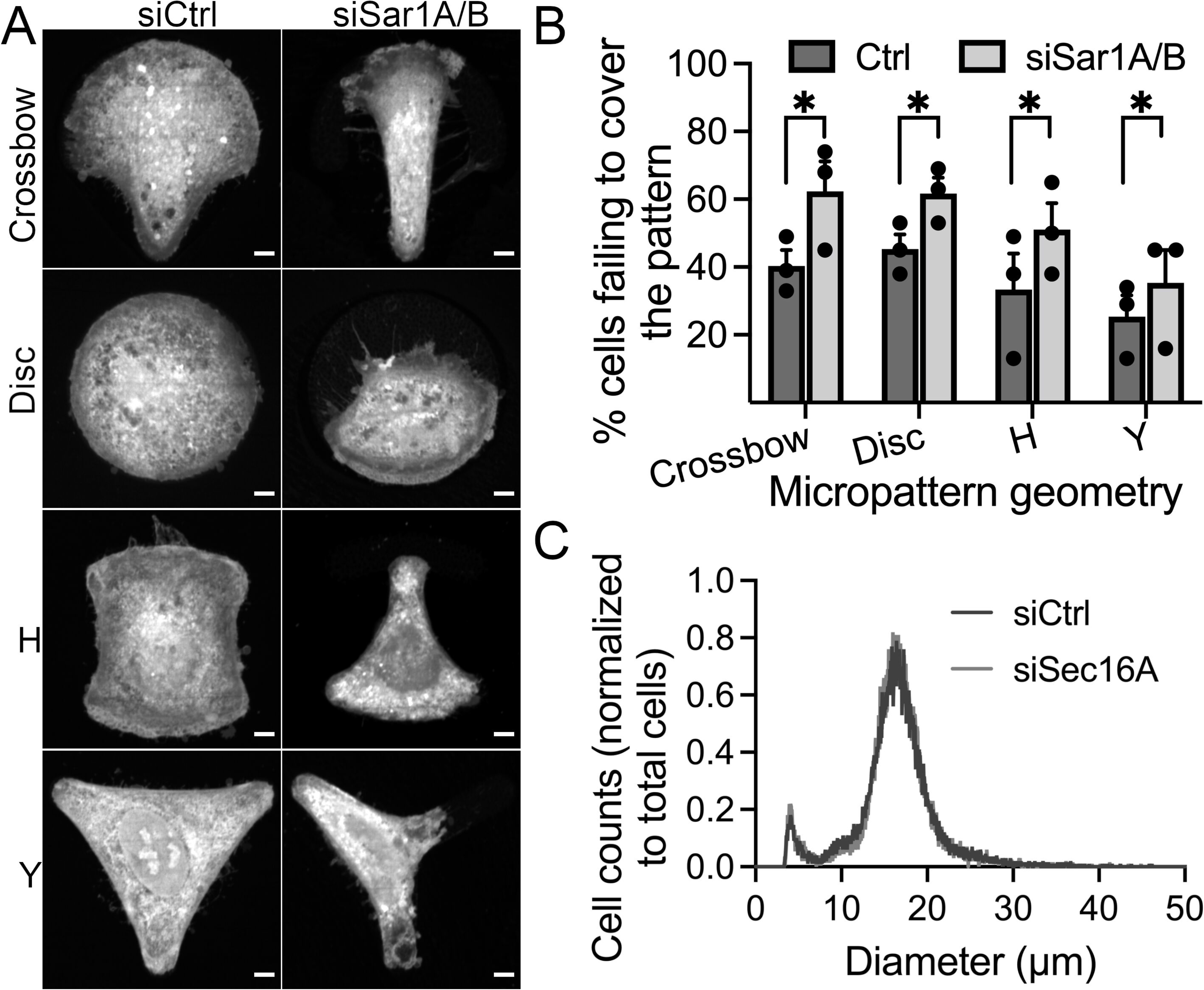
Dysfunctional ERES affects ability of cells to adapt to mechanical tension. (A) Representative immunofluorescence images of HeLa cells cultured on micropatterned surface of multiple geometries. Cells were transfected with 10 nM non-targeting control (siCtrl) or co-transfected with siRNAs targeting Sar1A and Sar1B. After 48 h, cells were trypsinized and 80,000 cells were seeded on micropatterned chips. Cells were allowed to attach for 3 h, stained with CellMask for 5 min and processed for microscopy. Scale bars: 5 µm (B) Quantification graph showing the percentage of cells failing to entirely cover the different geometries (described in A). Data derived from at least 100 cells from a set of three independent experiments. Asterisks (*) denote statistical significance (p-value < 0.05; chi-squared test). (C) Quantification graph shows the diameter of cells in non-targeting control (siCtrl) or siRNA targeting Sec16A (siSec16A). After trypsinization and resuspension of cells in culture medium, the diameter of cells was determined using the CASY cell counter. A total of 23,450 and 13,300 cells were counted for siCtrl and siSec16A, respectively.

### Perturbation of Rac1 activation impairs the early secretory pathway

We next asked whether Rac1 also signals to ERES in the absence of mechanical strain. For this purpose, we silenced Rac1 expression using siRNA in HeLa cells and assessed the number of ERES. We found that downregulation of Rac1 significantly reduced the ERES count (**Fig.3A-B**). We verified the depletion of Rac1 by Western blotting (**Fig. S3A**). Since Rac1 is required to maintain the cytoskeleton, we also measured the area covered by Rac1 knockdown cells and found no significant differences compared to control cells (**Fig.S3B**). The effect of Rac1 depletion on ERES could be rescued by introducing a siRNA resistant version of Rac1 in the cells (**Fig. S3C-E**). To further verify the results from the knockdown experiments, we treated HeLa cells with the Rac1 inhibitor NSC23766 for 4 hours, fixed and processed for quantification of ERES by immunofluorescence and confocal microscopy. In agreement with the results (**Fig.3A-B**) from knockdown experiments, pharmacological inhibition of Rac1 also led to a remarkable decrease in ERES (**Fig.3C-D**). Similar results were obtained in breast cancer MDA-MB-231 cells and prostate cancer PC3 cells (**Fig.S3F- G**), indicating that the effect is not cell type specific. Notably, ERES number in Rac1- knockout PC3 cells was not affected by treatment with NSC23766, thus ruling out that the observed effects on this inhibitor on ERES are due to an off-target effect (**Fig. S3H-J**). Because NSC23766 is a reversible inhibitor, we also checked whether the effect of Rac1 inhibition on ERES subsides following inhibitor washout. Indeed, the number of ERES almost completely recover after 2 h of NSC23766 washout (**Fig.3C-D**).

**Figure 3.**
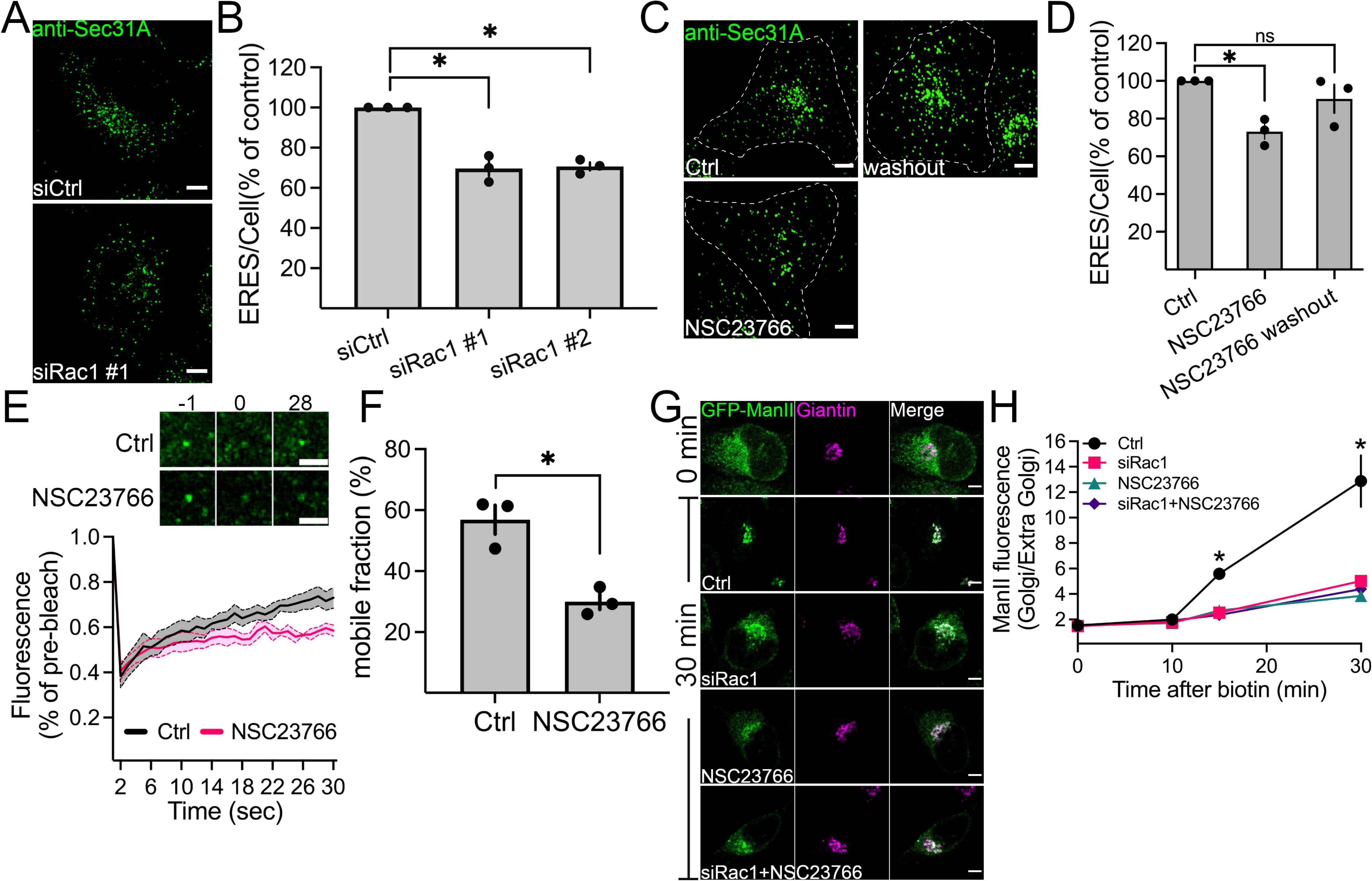
Perturbation of Rac1 activity affects ERES and ER-export. **(A)** HeLa cells were transfected with 10 nM non-targeting control siRNA (siCtrl) or siRNAs targeting Rac1 (siRac1). After 72 h, cells were fixed and processed for immunostaining against Sec31A to label ERES. Representative confocal microscopy images are shown. **(B)** Quantification graph shows the number of ERES/cell in cells transfected with control siRNA (siCtrl) or siRNA against Rac1 (two siRNAs #1 and #2). Values are expressed as % of siCtrl. From three independent experiments, with >30 cells per condition were counted. **(C)** Representative immunofluorescence images showing ERES in HeLa cells treated with DMSO (Ctrl), Rac1 inhibitor NSC23766 (50 µM, 4 h), or NSC23766 washout (50 µM for 4h, then washout for 2h). **(D)** Graph shows the number of ERES per cell (displayed as % of Ctrl) derived from at least 30 cells per condition from three experiments. **(E)** Fluorescence recovery after photobleaching (FRAP) of GFP-Sec16 marked ERES in HeLa cells. Images show an individual ERES before (-1), immediately after (0) and 28 sec after photobleaching. The graph illustrates FRAP analysis of individual ERES from 23 (for control) and 24 (for NSC23766) regions in 3 experiments. **(F)** Quantification of the mobile fraction of GFP-Sec16 in control and Rac1 inhibited cells. **(G)** The rate of ER-export was monitored using GFP-ManII-RUSH in HeLa cells after perturbation of Rac1 with siRNA (siRac1), or Rac1 inhibitor NSC23766, or a combination of both. Representative images show GFP-ManII-RUSH distribution in HeLa cells at indicated time points. **(H)** Quantification shows ratio of ManII fluorescence intensity within Golgi to outside Golgi region after addition of biotin at indicated time points. ManII intensity were measured in at least 30 cells in each experiment. Three independent experiments were performed. Scale bars in all immunofluorescence images are 5 µm. In all graphs, asterisks (*) indicate statistical significance (p-value < 0.05, Student’s unpaired *t* test), whereas ns indicates non- significant. Error bars represent SEM.

To determine the effect of Rac1 on ERES dynamics in living cells, we performed fluorescence recovery after photobleaching (FRAP) in cells expressing GFP-tagged Sec16A. We have previously used this strategy to uncover effects on ERES biogenesis and maintenance (Farhan *et al*., 2010; Tillmann *et al*, 2015). As shown in **Fig.3E-F**, inhibition of Rac1 reduced both the fluorescence recovery, resulting in a reduced mobile fraction of GFP- Sec16 after compared to control cells.

We wanted to further investigate whether the observed effect of perturbed Rac1 activity on ERES translates to altered ER-to-Golgi transport. We performed RUSH experiments with ManII as a cargo and noted a substantial delay in the rate of ManII arrival at the Golgi upon knockdown or inhibition of Rac1 (**Fig.3G-H**). Taken together, the results presented so far clearly establish an important role for Rac1 in regulating the ERES and the early secretory pathway.

### Manipulation of Rac1 activity at the ER reduces ERES number

Recent studies have uncovered that multiple and distinct subcellular pools of Rac1 regulate different cellular functions (Phuyal & Farhan, 2019). However, any presence of functionally active Rac1 at the ER remains unexplored. Therefore, we wondered whether Rac1 localizes to the ER. Because Rac1 antibodies do not work in immunofluorescence, we generated HeLa cells stably expressing mCherry-tagged Rac1. These cells were transiently transfected with GFP-tagged ER-resident protein Sec61β to label the ER. Using live cell imaging, we monitored for any co-occurrence of Rac1 and Sec61β. Because the majority of Rac1 is cytosolic, which could mask a subtle pool of this GTPase at the ER, we used SRRF imaging (Gustafsson *et al*, 2016). The localization of Rac1 at the ER might be transient and therefore difficult to detect, because Rac1 predominantly localizes to the plasma membrane and the cytosol. Using SRRF imaging, we detected a transient pool of Rac1 puncta co-occurring with Sec61β (**Fig.4A-B, movieS3**). To substantiate these findings, we generated HeLa cells in which the endogenous copy of Rac1 was tagged at the N-terminus with GFP using CRISPR/Cas. Using SRRF imaging we noted that endogenous Rac1 is detectable on the nuclear rim, which is a possible indication of its presence at the ER (**Fig.4C**) and we noticed the presence of puncta, which co-stained for the ERES marker Sec31A (**Fig.4D**). Thus, we provide the first evidence for the presence of Rac1 at the ER and at ERES.

**Figure 4.**
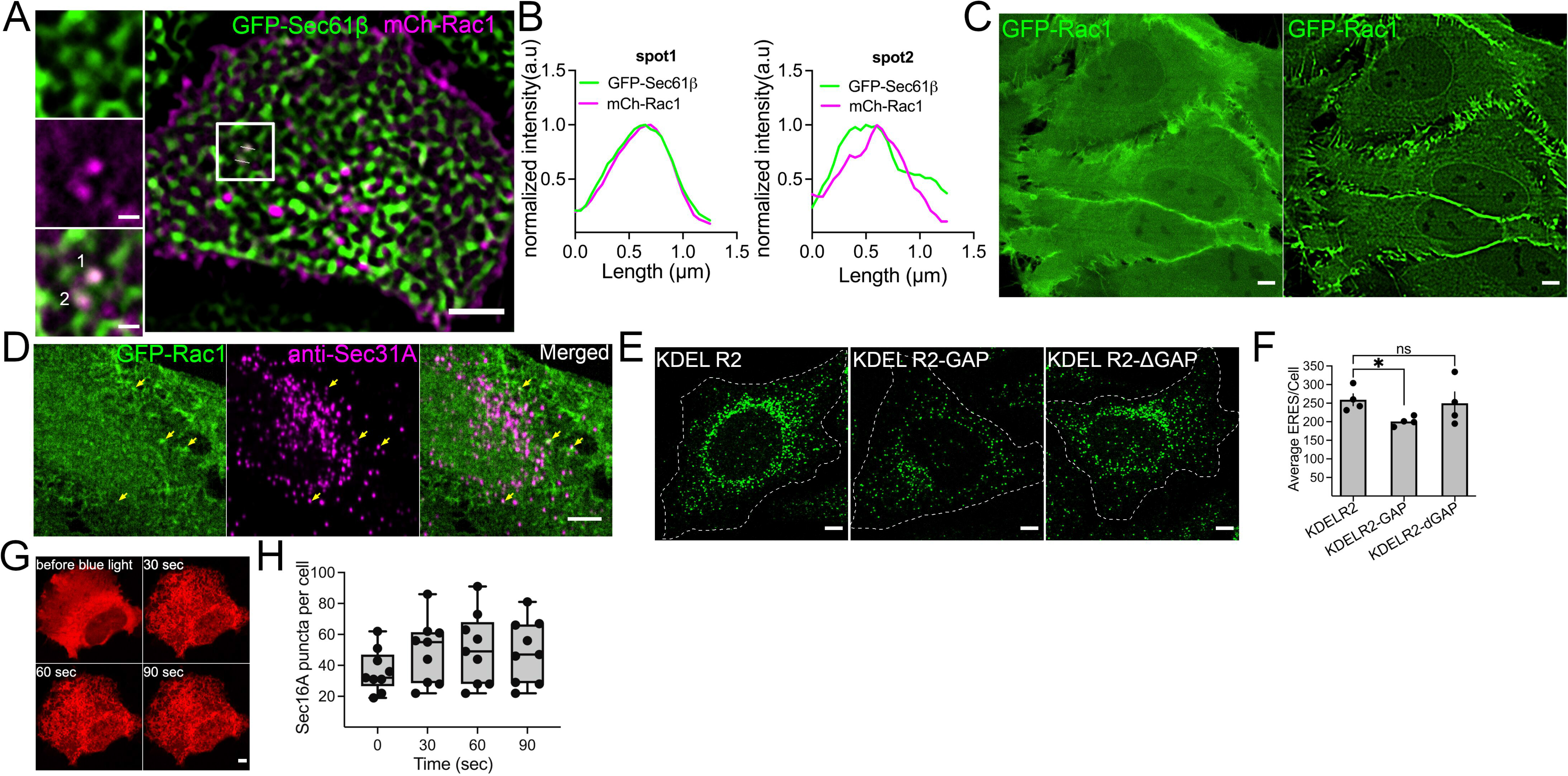
Manipulation of ER localized Rac1 affects ERES. (A) Co-occurrence of mCherry-Rac1 and GFP-Sec61β at the endoplasmic reticulum (ER) in HeLa cells stably expressing mCherry-Rac1 and transiently expressing GFP-Sec61 β as monitored by confocal microscopy. Images were processed using SRRF algorithm to improve the spatial resolution. Scale bar: 5 µm. See also movieS3. (B) Line profile of two marked spots from (A) showing the overlap between mCherry-Rac1 and GFP-Sec61β signal at the ER. (C) Side-by-side comparison of cells with endogenously tagged GFP-Rac1. The image on the left shows a confocal image and the image on the right was performed using SRRF. Scale bar: 5 µm (D) Cells with endogenous GFP-Rac1 were fixed and stained against Sec31A to label ERES. Arrows indicate Rac1-puncta colocalizing with ERES. Scale bar: 5 µm (E) Representative images showing Sec31A marked ERES in HeLa cells transfected with indicated plasmids. HeLa cells were transfected with the plasmids 16 h before immunolabelling with Sec31A. Scale bar: 5 µm. (F) Quantification of ERES number in HeLa cells from the experiment described in C. Between 90-100 cells were quantified from four experiments. Average ERES per cell ± SEM are shown. * = P-value < 0.05, ns = not significant (G) Images show distribution of Rac1 specific GEF TIAM in HeLa cells following optogenetic recruitment at indicated time points. CIBN-Sec61β was used to recruit Cry2- TIAM to the ER with 488 nm laser. Scale bar: 5 µm. (H) Graph shows the number of SNAP-Sec16A marked ERES in HeLa cells before (0) and after (30, 60 and 90 seconds) recruitment of TIAM to the ER. Raw data distribution and min- max range are shown. Number of cells = 9, number of experiments = 2.

This observation prompted us to test whether the ER-localized pool of Rac1 could be selectively manipulated and whether this would lead to altered ERES. To selectively inactivate Rac1 at the ER, we generated a plasmid expressing the GAP-domain of Rac1- specific GAP β2-Chimaerin fused to the KDEL receptor 2 (KDELR2) for ER targeting (hereafter KDELR2-GAP). As a control, we created catalytically inactive GAP fused to KDELR2 by mutating the R363 to alanine (hereafter KDELR2-i”GAP). We also included KDELR2 as an additional control in our experiments. We overexpressed these plasmids in HeLa cells and quantified the effect on ERES. While KDELR2 or KDELR2-i”GAP overexpressing cells had similar number of ERES, cells overexpressing KDELR2-GAP exhibited a significant ERES reduction (**Fig.4E&F**). These results are similar to those obtained with the knockdown or the pharmacological inhibition of Rac1, which affect every subcellular pool of Rac1. However, the KDELR2-GAP only acts at the ER and thus these results point towards a functionally active pool of Rac1 at the ER possibly regulating the ERES.

We next asked whether increasing Rac1 activation at the ER is sufficient to regulate ERES. To this end, we took advantage of CIBN-CRY2 based light inducible dimerization system (Kennedy *et al*, 2010) to recruit the Rac1 specific GEF TIAM1 to the ER (**Fig.4G**). This approach of generating spatially restricted active Rac1 has previously been used to activate Rac1 at the leading edge in cells (de Beco *et al*, 2018). As presented in **Fig.4G**, cytosolic TIAM was successfully recruited to the ER by CIBN fused Sec61β. With this approach we activated Rac1 locally at the ER in living HeLa cells and examined changes in ERES using SNAP-tagged Sec16. We noted an overall increase in ERES over time (**Fig.4H**) further strengthening the notion of an involvement of active Rac1 in signaling to ERES.

### Effect of Rac1 on the early secretory pathway is actin independent

We next aimed to unravel the underlying molecular details of Rac1 regulated ER-export. Rac1 is pivotal in regulating actin dynamics and actin assembly has previously been observed at the ER (Wales *et al*, 2016). Therefore, we asked whether actin mediates the observed effect of Rac1 on ERES. To test this possibility, we treated HeLa cells with different concentrations of two actin disrupting agents cytochalasin D (CytoD) and latrunculin A (LatA) for 15 and 30 min. We then quantified the effect of actin disruption on ERES and ER-to-Golgi transport. Neither CytoD nor LatA treatment reduced ERES (**Fig.5A-B**) or affected the rate of the ER-to-Golgi transport (**Fig.5C**) as Rac1 did. It should be mentioned that in live cell imaging experiments, using the GFP-tagged actin probe actin-chromobody, we verified that both drugs worked as anticipated (**Fig. S4A**). To perform a more targeted actin perturbation, we silenced the expression of inverted formin 2 (INF2) (**Fig.S4B-C**), which was previously reported to regulate actin dynamics locally at the ER (Wales *et al*., 2016). We determined the number of ERES (**Fig.5E-F)** or the kinetics of ER-to-Golgi transport (**Fig.5G-H**) in INF2 depleted cells but were unable to detect any appreciable changes compared to the control.

**Figure 5.**
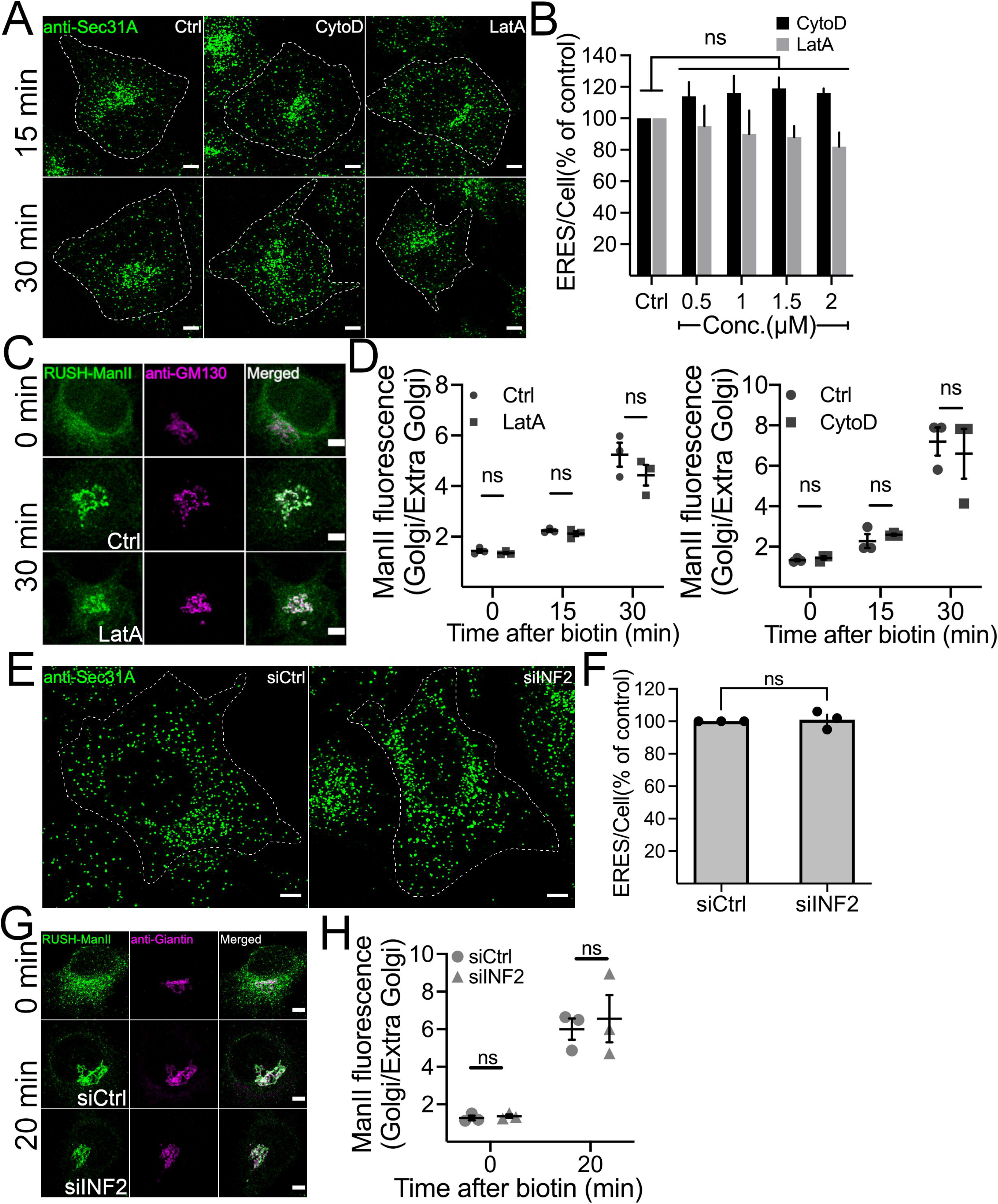
Effect of actin disruption on ERES and ER-to-Golgi transport. (**A** and **B**) Representative immunofluorescence images showing Sec31A labelled ERES in HeLa cells treated with DMSO (Ctrl), 0.5 µM cytochalasin D (CytoD) and 0.5 µM latrunculin A (LatA) for 15 and 30 min (A). Scale bar: 5 µm. The number of ERES per cell in HeLa cells treated with different doses of CytoD and LatA for 30 min are displayed as % of control ± SEM (90 -100 cells from 3 experiments) (B). (**C**) Distribution of ManII-RUSH in HeLa cells. Cells were pre-treated with DMSO (Ctrl) or 1 µM LatA for 15 min prior to addition of biotin to initiate ManII-trafficking. Golgi arrival for ManII-RUSH was monitored for 30 min. Scale bar: 5 µm. (**D**) Graph showing ratio of ManII within Golgi to outside Golgi. Values were derived from HeLa cells treated with LatA or CytoD as described in C. Error bars represent SEM. At least 30 cells per condition were used for quantification in each experiment (n = 3). (**E** and **F**) siRNA mediated depletion of INF2 does not phenocopy Rac1 effect. HeLa cells were transfected with non-targeting control siRNA (siCtrl) or siRNA against INF2 (siINF2) for 72 h before processing cells immunostaining with Sec31A to quantify ERES. Representative images showing Sec31A-labelled ERES in HeLa cells (E), and quantification of ERES per cell presented as % of control (F). Scale bar in E is 5 µm, error bars in F represent SEM. (**G** and **H**) Knockdown of INF2 does not alter the kinetics of ER-to-Golgi transport. HeLa cells stably expressing ManII-RUSH were transfected with siRNAs as described in E, and the ER-to-Golgi transport of ManII-RUSH was followed by confocal microscopy (G). Representative images for indicated time points are shown. Scale bars, 5 µm. The ratio of ManII-RUSH within the Golgi to outside the Golgi was calculated for the indicated time points (H). Error bars represent SEM. In all graphs, ns = not significant.

Based on these data, we conclude that the effect of Rac1 on the early secretory pathway is actin independent.

### Rac1 interacts with the COPII subunit Sar1 and stimulates vesicle budding

Because we observed functionally active Rac1 at the ER (**Fig. 4**), we wondered whether Rac1 directly signals to the constituents of the ER-export machinery. We first visualized mCherry-tagged Rac1 together with GFP-tagged Sec16 in HeLa cells. Interestingly, we observed transient co-occurrence of Rac1 with Sec16 (**Fig.6A, movieS4**) further suggesting a direct link between Rac1 and ERES. At ERES, recruitment of the small GTPase Sar1 is an early step during the process of COPII assembly. Therefore, we investigated whether Rac1 may interact with Sar1. Because the localization of Rac1 to ERES was very transient (**movie S4)**, we opted for biomolecular fluorescence complementation (BiFC) to capture transient Rac1- Sar1 interactions. We co-expressed YPF(N)-tagged Rac1 and YFP(C)-tagged Sar1 in HeLa cells and observed that many of the Rac1-Sar1 complexes formed puncta that colocalized with Sec31A, indicating that they are ERES (**Fig.6B-C**). Moreover, we were also able to coimmunoprecipitate endogenous Sar1 with GFP-tagged Rac1 in HeLa cells (**Fig. 6D**) further supporting results from the BiFC assay (**Fig.6B**). We also used BiFC to probe for Rac1 and Sec16 interaction and were unable to detect any interaction (**Fig.S5**). Together these findings indicate that Rac1 may regulate secretion via its interaction with Sar1.

**Figure 6.**
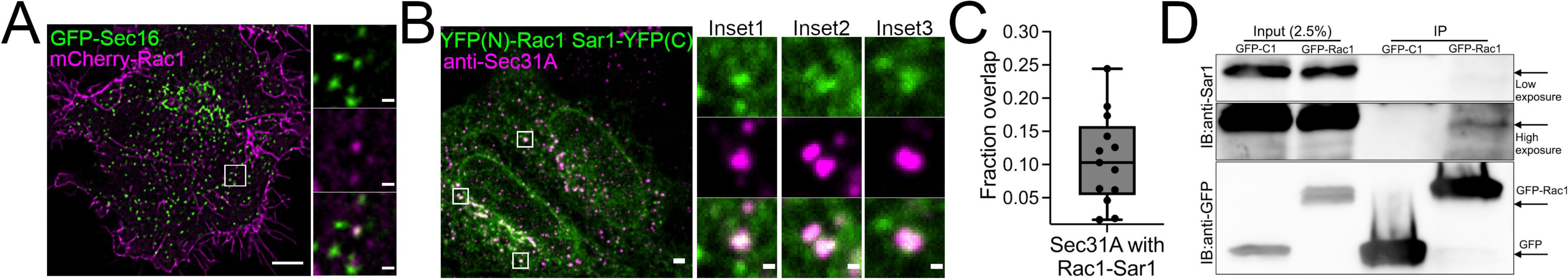
Rac1 regulates ERES and ER-export by dimerizing with Sar1. (A) Co-localization of mCherry-Rac1 and GFP-Sec16A in HeLa cells as monitored by SRRF live cell imaging. See also movieS4. (B) Representative immunofluorescence images of HeLa cells co-transfected with YFP(N)- Rac1 and Sar1-YFP(C) and immunostained with Sec31A. Scale bars: 5 µm. Rac1 and Sar1 form complexes at ERES in HeLa cells, which partially overlap with Sec31A. (C) Box plot showing the fraction of ERES that overlap with the Rac1-Sar1 complex (number of cells = 24, number of experiments = 4). Individual data points shown in the graph represent number of images. (D) GFP-tagged Rac1 was immunoprecipitated by GFP-trap and the fraction was immunoblotted for endogenous Sar1. Representative immunoblot showing co-immunoprecipitation of Sar1 by Rac1.

We next explored the Rac1-Sar1 interaction using *in silico* modeling, which suggested that Rac1 could cover the GTP binding site in Sar1 (**Fig.7A-B**). In accordance with this, the *in silico* model predicts that Rac1 binding to Sar1 might block Sec23 (Sar1 GAP) from accessing Sar1, due to a steric clash between Sec23 and Rac1 (black circle in **Fig.7C**). On the contrary, our model predicts that the Sar1-Rac1 dimer can interact with the Sar1 GEF Sec12 (**Fig.7D**). To validate this prediction, we performed co-immunoprecipitation experiments. As shown in **Figure 7E**, immunoprecipitated GFP-tagged Rac1 brought down endogenous Sar1 and Sec12, supporting the notion of the existence of a ternary complex and increasing the level of confidence in our *in silico* model.

**Figure 7.**
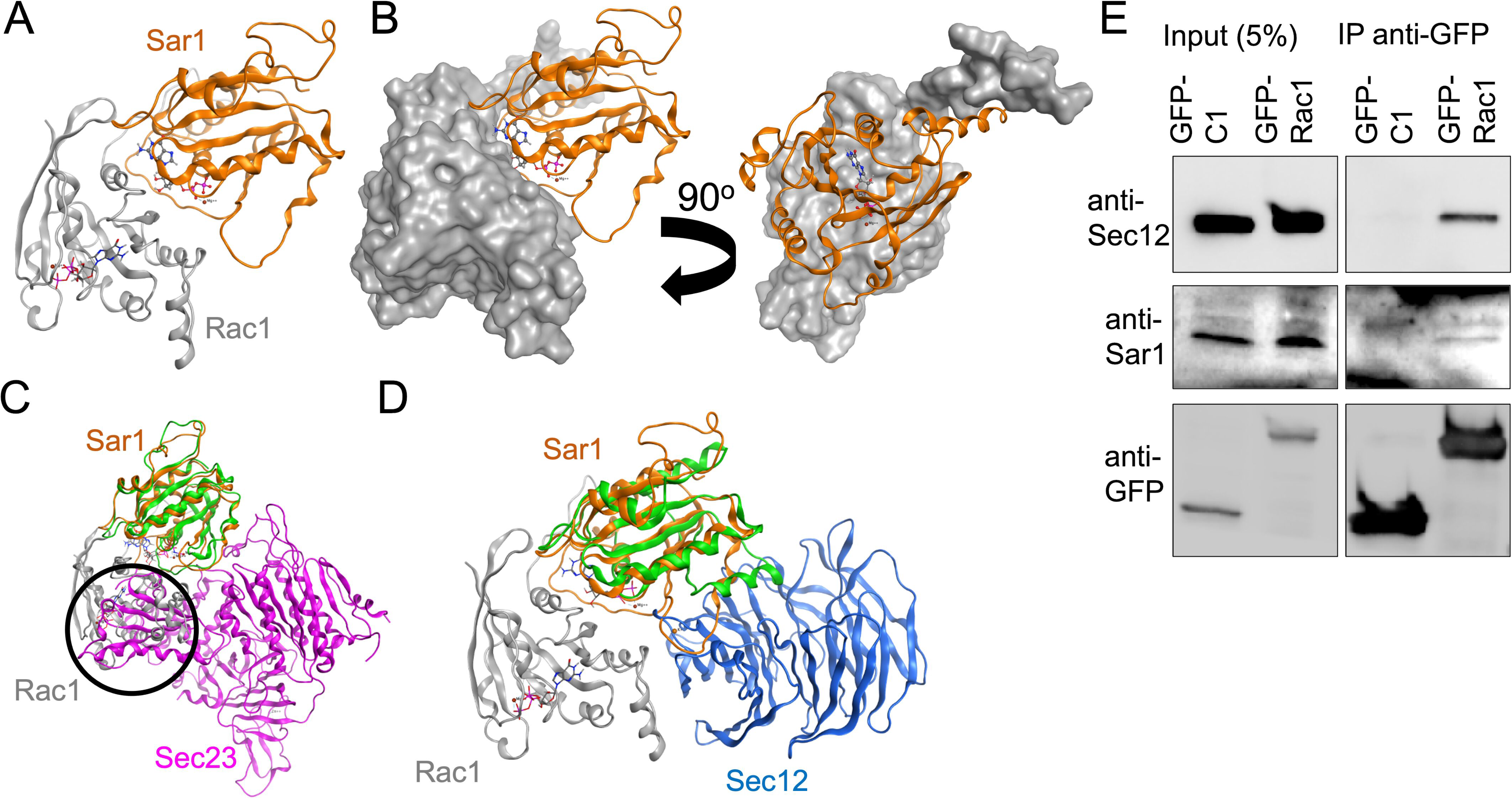
In silico modeling predicts a Rac1-Sar1-Sec12 ternary complex and its experimental validation. (**A&B**) *In silico* model showing the complex between Rac1 (grey) and Sar1 (gold) and how Rac1 covers the GTP binding site of Sar1. GTP of Rac1 in stick model, and of Sar1 in ball-and-stick. (C) *In silico* model showing the steric clash between Rac1 and Sec23 (encircled), obtained by superposing the Sar1 units of the Sar1-Sec23 (green-purple) heterodimer onto the Sar1-Rac1 heterodimer (gold-grey). (D) *In silico* model showing that the Sar1-Rac1 heterodimer can interact with Sec12 by superposing the Sar1 units of Sar1-Rac1 (gold-grey) and Sar1-Sec12 heterodimers (green-blue). (E) Cells expressing GFP-Rac1 were lysed followed by immunoprecipitation against GFP and immunoblotting against endogenous Sar1 and Sec12. Blotting against GFP was performed to determine efficiency of the IP.

Because Rac1 has a positive effect on ERES structure and function, we reasoned that Rac1 ought to exert a positive effect on Sar1. This can occur either as a consequence of enhanced Sar1 activation by Sec12, or because of reduced inactivation by Sec23. Our *in silico* model predicts that Sec23 is not capable of binding the Sar1-Rac1 dimer. On the other hand, our combined experimental and computational data indicate that Sec12 can bind the complex. Therefore, we next investigated whether Rac1 affects the activation state of Sar1. To this end, we incubated microsomes with recombinant Sar1 followed by pulldown with an antibody directed to the active form of Sar1. Microsomes contain Sec12, the GEF for Sar1, and indeed we observed Sar1 activation on microsomes (**Fig.8A**). When we included recombinant Rac1 in the assay, we obtained higher levels of active Sar1 (**Fig.8A**).

**Figure 8.**
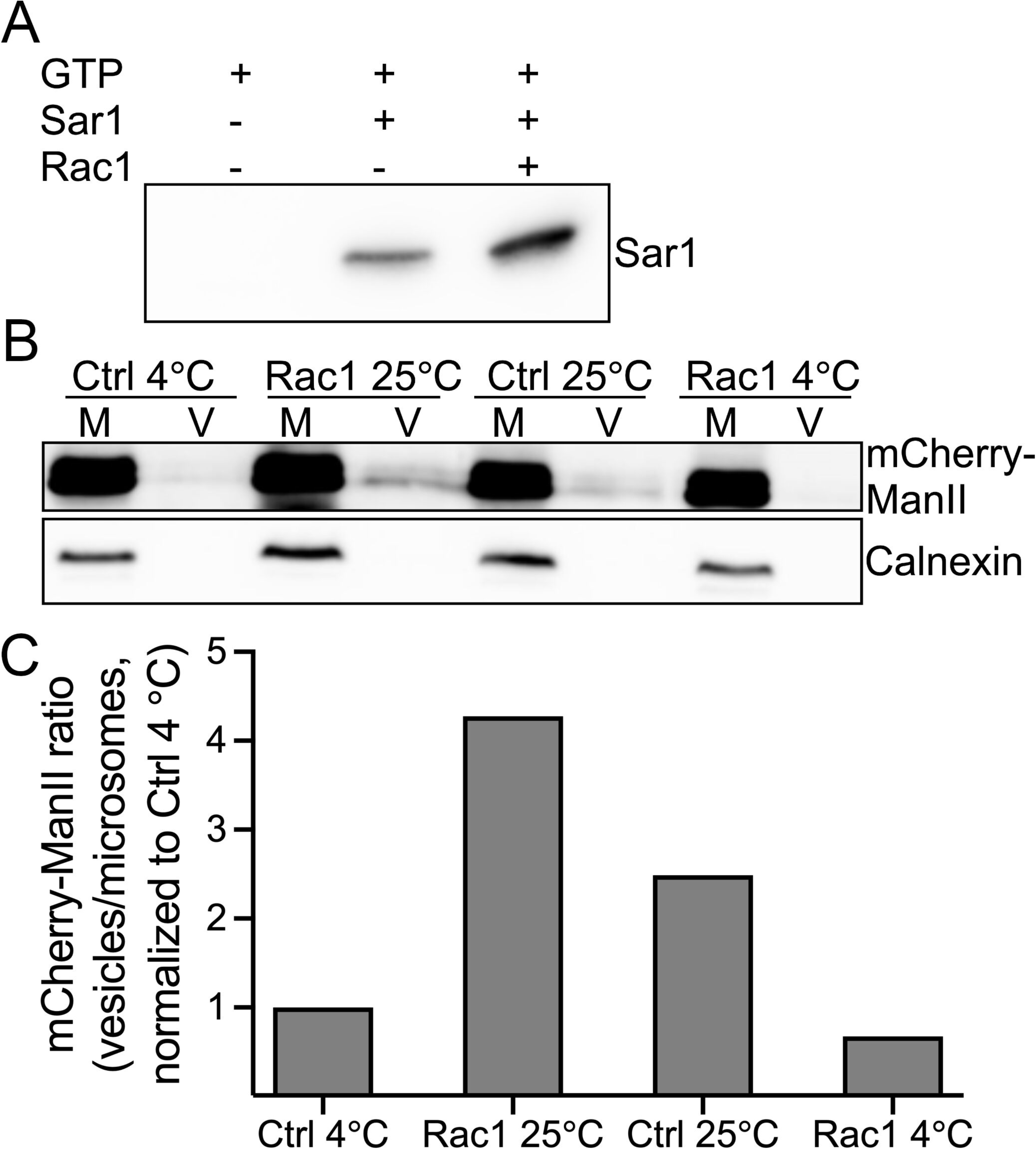
Rac1 stimulates the formation of active Sar1 and COPII. (A) Microsomes isolated from HeLa cells were incubated as indicated. Active Sar1 pulldown was then carried out, followed by immunoblotting against Sar1. (B) Microsomes (M) from HeLa cells stably expressing mCherry-ManII-RUSH were incubated with ATP and an ATP-regenerating system, GTP, biotin and cytosol in the presence or absence of recombinant Rac1 for 30 min at 25°C. COPII vesicles (V) were pelleted by ultracentrifugation (100,000 xg) and immunoblotted for anti-mCherry to detect ManII-RUSH and anti-calnexin (ER marker). (C) Quantification of two vesicle budding assays. All values were normalized to 4 C Ctrl condition.

Active Sar1 promotes COPII assembly and thereby the formation of carriers from the ER. To determine whether recombinant Rac1 stimulates carrier formation, we used an *in vitro* COPII vesicle budding assay (Farhan *et al*., 2010; Kim *et al*, 2005). We used microsomes from HeLa cells stably expressing the RUSH cargo ManII. In the absence of biotin, ManII is only found in the ER. Microsomes were incubated with cytosol from HeLa cells, biotin, recombinant Rac1, GTP and ATP regeneration system (see materials and methods for details). Vesicles and microsomes were separated by differential centrifugation. Performing the assay in the presence of Rac1 increased the amounts of mCherry-ManII in the vesicle fraction compared to the fraction without exogenous Rac1 (**Fig.8B-C**). This data, together with the result from the active Sar1 pulldown (**Fig.8A**), suggest that Rac1 stimulates cargo exit from the ER through its interaction and positive modulation of the COPII subunit Sar1.

## Discussion

In the current work, we showed that Rac1 regulates the stretch-induced increase in ERES and ER-to-Golgi transport. Rac1 forms a complex with Sar1 and Sec12 and increases Sar1 activation. Because Sar1 is a major regulator of ER-export, we suggest that the Rac1 regulates secretion through binding to Sar1. Mounting evidence suggests that small GTPases form dimers, consequently affecting their biological function. For instance, dimerization of Arf1 (Diestelkoetter-Bachert *et al*, 2020) and Sar1 (Hariri *et al*, 2014) was shown to regulate biogenesis of COPI vesicles and scission of COPII vesicles, respectively. Reportedly, Rac1 homodimerization regulates its intrinsic GTP hydrolyzing ability (Zhang *et al*, 2001). In contrast to homodimerization, only scant information is available on heterodimerization of small GTPases. Hereby, we propose that Rac1 and Sar1 form such a heterodimer based on co-immunoprecipitation and BiFC data. All other downstream biological effects of Rac1 on ERES are compatible with the positive effect of Rac1 towards Sar1. This raises the question of how does the Rac1-Sar1 dimer positively regulate ER-export? The fact that we observe a ternary complex of Rac1, Sar1 and Sec12 suggests that Rac1 might promote Sar1 activation. This is supported by our finding of more Sar1-GTP formed on microsomes in the presence of Rac1. In addition, our in silico data suggest that the Rac1-Sar1 complex does not accommodate Sec23. A possible alternative interpretation of our data is that Rac1-Sar1 heterodimerization prevents formation of Sar1 homodimers. Sar1 homodimerization was proposed previously to promote vesicle scission (Hariri et al, 2014), and therefore the heterodimerization with Rac1 might delay vesicle scission, thereby extending the time for cargo loading and the formation of larger carriers. At the moment we can only speculate about the existence of such carriers. In support of this speculation is our live imaging experiment where we observed that stretched cells exhibited more tubular ERES. However, we point out that the topic of whether ERES give rise to vesicles tubules or tunnels is a controversially discussed topic (Phuyal & Farhan, 2021) and further explorations along this line are certainly beyond the scope of the current work.

Another surprising observation was the identification of a very transient and small pool of Rac1 at the ER and at ERES. The high level of cytosolic Rac1 in any given cell type might have precluded the identification of this subcellular Rac1 pool with conventional confocal microscopy techniques. The ER pool of Rac1 is biologically active and is unlikely representing nascent biosynthetic Rac1. This is supported by the data showing that localized inactivation of Rac1 at the ER (with an ER-targeted GAP) reduced the number of ERES. In addition, Rac1 formed complexes with Sar1 at ERES. The observation of Rac1 at the ER is in line with a previous finding (Woroniuk *et al*, 2018), which documented Rac1 signaling activity at the perinuclear region, a membrane system that is continuous with the ER. Reportedly, the C-terminal fusion of CAAX motif of Rac1 to the green fluorescent protein GFP was sufficient for targeting GFP to the ER (Choy *et al*, 1999) further highlighting the intrinsic ability of Rac1 to associate with the ER membrane.

Our work establishes for the first time a mechanistic link between mechanotransduction and the regulation of ERES and demonstrates an important role of ERES for proper cellular adaptation in mechanically challenged cells. Previous research has established the regulation of ERES by intracellular stimuli (e.g. mitosis or unfolded proteins in the ER) or by extracellular chemical stimuli (e.g. nutrients and mitogens). Our work completes the picture of the regulation of ERES by mechanical stimuli. This will pave the way for future investigations on the role of other mechanical stimuli such as stiffness, compression or shear stress. The mechanisms by which Rac1 activity is regulated during mechanotransduction are poorly understood, although studies have proposed a role for mechanosensitive channel and focal adhesion molecules at the cell surface (Arya *et al*, 2020; Bae *et al*, 2014). Future work is needed to characterize the receptors or sensors at the cell surface that signal to Rac1.

Finally, our work has implications for the regulation of proteostasis in response to changes in cell size. We noticed that the absolute number of ERES is higher in cells forced to grow on larger micropatterns. The same was true when we expanded the cell area using acute stretching. This shows that cells are capable of adapting their secretory capacity when they expand in size. In fact, a recent pre-print showed that different organelles undergo changes dependent on the cell size (Lanz *et al*, 2021). Future work could focus on studying how cell size affects proteostasis by controlling the rate of protein synthesis, secretion and degradation.

## Materials and Methods

### Cell culture and treatment

HeLa cells were maintained in DMEM (Lonza, #12-604F) supplemented with 10% FCS (Life Technologies, #10500064), 50 units/ml penicillin and 50 µg/ml streptomycin (Lonza, #DE17- 603E), in a humidified 5% CO2 atmosphere at 37 °C. PC3 cells and MDA-MB-231 cells were cultured in RPMI medium with supplements as for HeLa cells. CRISPR Rac1 knockout PC3 cells were extensively characterized previously (Baker *et al*, 2020).

For all Rac1 inhibition experiments, cells were treated with 50 µM NSC23766 or 0.1% (v/v) DMSO in complete medium for 4 h. For actin disruption experiments, cells were treated with 0.5, 1, 1.5 and 2 µM cytochalasin D or latrunculin A in complete medium for 15 and 30 min. For RUSH assays, cells were preincubated with 1 µM cytochalasin D or latrunculin A for 15 min before starting the cargo release with biotin.

siRNA oligos were delivered to the cells using HiPerFect transfection reagent. Briefly, 10 - 20 nM siRNA duplexes were complexed with the transfection reagent in 100 µl serum and antibiotics free cell culture medium, incubated at room temperature for 5 min and added to the cells immediately after seeding for experiments. Knockdown efficiency after 72 h was measured to check that the levels of desired proteins were reduced. For Rac1 knockdown experiments, the siGenome non-targeting siRNA pool from Dharmacon were used as control. The AllStars negative control siRNA from Qiagen was used as control for INF2 knockdown experiments.

All plasmid transfections were carried out using Fugene 6 (used at 3 µl per µg DNA) transfection reagent following the manufacturer’s protocol. Cells were transfected with 750 ng of plasmid DNA. Unless otherwise stated, all listed concentrations of siRNAs and chemicals are final concentrations.

### Immunofluorescence staining

Cells were fixed with 4% paraformaldehyde for 12 min, washed three times with PBS, quenched for 5 min with 50 mM NH4Cl, and washed again twice with PBS. After permeabilization with 0.2% (v/v) triton X-100 for 4 min, cells were washed twice with PBS, and blocked for 30 min in 1% (wt/vol) BSA. Finally, cells were incubated for 1 h with the primary antibody, washed thrice with PBS, incubated for 1 h with secondary antibody and mounted with antifading Polyvinyl alcohol mounting medium with DABCO® (Sigma, #10981) after washing three times with PBS. All incubation steps were carried out at room temperature, triton X-100 and BSA dilutions were prepared in PBS, and each PBS washing steps were 5 min on a shaker.

### FRAP microscopy

HeLa cells were seeded on 35 mm MatTek dishes (MatTek corporation, #P35G-1.5-20-C) and incubated overnight. Cells were then transfected with 0.75 µg of GFP-Sec16 plasmid and incubated overnight. Prior to FRAP experiments, cells were treated with DMSO or with Rac1 inhibitor NSC23766 for 4 h. The experiment was performed on a Zeiss LSM700 confocal microscope equipped with a 63× oil-immersion objective (Plan-Apochromat 63x/1.40 NA Ph3 M27) at 37 °C. Images were acquired using Zeiss Zen software. A pre-bleach image was acquired before bleaching individual ERES at 100% laser intensity (10 iterations) and subsequent image acquisition at one image per second.

FRAP analysis was carried out as described previously(Centonze *et al*., 2019b; Farhan *et al*., 2010).

### Preparation of polydimethylsiloxane (PDMS) membranes

Stretchable PDMS (Sylgard Silicone Elastomer Kit, Dow Corning) membranes were prepared as previously described(Kosmalska *et al*, 2015). Briefly, a mix of 10:1 base to crosslinker ratio was spun on plastic wafers for 1 minute at 500 rpm and then cured overnight at 65 °C. Once polymerized, membranes were peeled off the wafers and assembled onto a metal ring for fibronectin coating, cell seeding and microscopy experiments.

### Mechanical stimulation of the cells and live cell imaging

Cell mechanical stimulation was done as previously described (Kosmalska *et al*., 2015). Briefly, a 150 µL droplet of a 10 µg/mL fibronectin solution (Sigma, #F0895) was deposited in the center of the membrane mounted in the ring. After overnight incubation at 4 °C, the fibronectin solution was rinsed. HeLa cells expressing GFP-Sec16A were then seeded on the fibronectin coated membranes and allowed to attach for up to 1 h. Then ring-containing membranes were mounted in the stretch system previously described(Kosmalska *et al*., 2015). Live-cells were subjected to a 7.5 % equibiaxial strain. Images of the cells were acquired with 500 ms interval frames, with a 60x objective (NIR Apo 60X/WD 2.8, Nikon) mounted in an inverted microscope (Nikon Eclipse Ti) with a spinning disk confocal unit (CSU-W1, Yokogawa), a Zyla sCMOS camera (Andor) and using the Micromanager software.

For Rac1-FRET assay and the RUSH assay, HeLa cells were also subjected to a 7.5 % equibiaxial strain. Images were acquired with a 60x objective (NIR Apo 60X/WD 2.8, Nikon) mounted in an upright epifluorescence microscope (NIR Apo 60X/WD 2.8, Nikon) and an Orca Flash 4.0 camera (Hamamatsu), using the Metamorph software.

### Live cell imaging and optogenetics

Live cell imaging of HeLa cells co-expressing mCherry-Rac1 and GFP-Sec61 or mCherry- Rac1 and GFP-Sec16 was performed on Andor Dragonfly spinning disk equipped with a Nikon Ti2E inverted optical microscope (60 × TIRF objective (Plan-APOCHROMAT 60 × /1.49 Oil) and an EMCCD camera (iXon Ultra 888, Andor). Cells were maintained at 37 °C in 5% CO2 in a heating chamber (Oko lab). SRRF-Stream mode in Fusion (version 2.1, Andor) was used for image acquisition with an additional 1.5 × magnification. Following imaging parameters were used. SRRF Frame count: 150-200, SRRF Radiality Magnification: 4×, SRRF Ring Radius: 1.4 px, SRRF Temporal Analsysis: Mean and SRRF FPN correction: 75 -100 frames.

For optogenetics, HeLa cells expressing CIBN-GFP-Sec61β, Cry2-TIAM-mCherry and SNAP-Sec16A were stimulated for 30 sec with a 488 nm laser.

### Retention using selective hooks (RUSH) assay

A step-by-step protocol for RUSH assay has been described elsewhere(Boncompain & Perez, 2012). We used HeLa cells stably expressing the Golgi resident protein Mannosidase II in RUSH system, and monitored its arrival at the Golgi for up to 30 minutes in our experiments.

### ERES, RUSH quantification and colocalization analysis

ERES and RUSH quantification, and colocalization analysis were all carried out using Fiji(Schindelin *et al*, 2012). Prior to analysis, background was subtracted from the immunofluorescence images using “subtract background” command in Fiji. To obtain ERES count in cells, images were thresholded (Otsu algorithm) and ERES were counted using “Analyze particle” command. Counts were then normalized to total cells.

In live cell imaging equibiaxial strain experiments, the number of GFP-Sec16 marked ERES in each frame were detected and counted using “Find maxima” in Fiji.

For quantification of RUSH experiments, ManII intensity in the Golgi was measured by drawing a ROI. To measure extra Golgi intensity of ManII, the ROI was then transferred outside the Golgi. Ratio of ManII at the Golgi to extra Golgi region was calculated for data visualization. The fraction overlap of Rac1-Sar1 complex with Sec31 was calculated using DiANA(Gilles *et al*, 2017) plugin in Fiji.

### Cell lysis, immunoprecipitation and immunoblotting

Cells were washed twice with ice-cold PBS, scraped off in lysis buffer (50 mm Tris-HCl, 300 mm NaCl, 1 mm EDTA, 0.5% Triton X-100, pH 7.4) supplemented with EDTA free proteinase and phosphatase inhibitor mixture and incubated in ice for 10 min. Cell lysates were then centrifuged at 20,000 xg for 10 min at 4 °C. The supernatant was collected and solubilized in loading buffer. Samples were run on 4-20% TGX gels (BioRad) and proteins were transferred to a nitrocellulose membrane using semidry transfer system. Once the transfer was complete, membranes were blocked (in 5% (wt/vol) milk in TBS with 0.1% Tween) and incubated first with the specified primary antibodies, washed and then with the HRP-conjugated secondary antibodies. Finally, blots were visualized using ECL clarity chemiluminescence reagent (BioRad, #1705061) on ChemiDoc (BioRad).

For immunoprecipitation experiments, cells were lysed in immunoprecipitation buffer (20 mM Tris-HCl, pH 7.4, 150 mM NaCl, 1 mM MgCl2, 10% glycerol, 0.5% NP-40) and GFP-trap (ChromoTek, #gta-100) was performed following manufacturer’s protocol.

### Active Sar1 pulldown assay

Sar1 activation assay kit (NewEast Biosciences, #81801) and the protocol provided with the kit was followed for active Sar1 pulldown from microsomes.

To isolate microsomes, HeLa cells were scraped in ice-cold PBS and pelleted by centrifugation at 500 ×g for 10 min. The pellet was resuspended in 400 µl of buffer containing 10 mM Hepes-KOH, pH 7.2, 250 mM sorbitol, 10 mM KOAc, and 1.5 mM MgOAc. Cells were homogenized by passing through a 25-gauge syringe for at least 30 times and centrifuged at 1,000 ×g for 10 min. The postnuclear supernatant was then centrifuged at 6,000 g for 10 min. The pellet was resuspended in 400 µl of resuspension buffer (20 mM Hepes-KOH, pH 7.2, 250 mM sorbitol, 140 mM KOAc, and 0.5 mM MgOAc) and centrifuged at 6,000 ×g for 10 min. Finally, the pellet was resuspended in 500 µl of resuspension buffer and used for active Sar1 pulldown.

To the microsomal fractions, 750 ng of recombinant Rac1 or Sar1 or both and GTP or GTPyS (100 µM final concentration) were added. The reaction mixture was then incubated at 30 °C with constant agitation for 30 min for GTP-loading prior to active Sar1 pulldown with anti-active Sar1 monoclonal antibody provided in the kit.

### Vesicle budding assay

We prepared microsomes and cytosol for vesicle budding assay following the previously published protocol from the lab(Farhan *et al*., 2010). Microsomes were isolated from HeLa cells stably expressing ManII-RUSH, whereas cytosol was prepared from HeLa cells.

For budding reactions, 30 µl of microsomes, 65 µl of cytosol, 40 µM biotin, 0.5 mM GTP and an ATP regeneration system (1 mM ATP, 25 mM creatine phosphate, and 0.3 mg/ml creatine kinase) was used. Vesicle budding reactions were carried out either at 25°C or on ice (control) for 30 min. In the Rac1- and Sar1- containing reactions, 500 ng recombinant proteins were added. The reaction was stopped by placing the tubes on ice. Finally, the reaction mixture was centrifuged at 100,000 g for 30 min to pellet the budded vesicles.

### Statistical analyses

All statistical analysis was performed in Microsoft Excel. Where appropriate, we performed a Student’s unpaired *t* test to evaluate statistical significance of the data. In the figures, data are presented as mean ± SEM from at least three independent experiments, unless otherwise indicated in the figure legends. In the figures, asteriks (*) denotes a P-value < 0.05.

### Rac1 GFP tagging

Rac1 was gene edited using a modified PITCh technology (Sakuma *et al*, 2016) adjusted for generating N-terminal fusion proteins (Lin *et al*, 2019). A Rac1 specific gRNA 5’ ACACTTGATGGCCTGCATCA was cloned into plasmid pX330-BbsI-PITCh (Addgene plasmid #127875) and transfected along with pN-PITCh-GFP-Rac1 into HeLa cells using JetPrime (Polyplus, Illkirch, France). Plasmid pN-PITCh-GFP-Rac1 contains a Puromycin resistance-T2A-eGFP cassette flanked by 50 bp of genomic RAC1 sequence flanking the CRSIPR-CAS9 induced DNA double strand break. This plasmid was constructed by HiFi- mediated in vitro recombination (NEB) of two PCR based fragments generated by amplifying the pN-PITCh-GFP (Addgene plasmid #127888) vector backbone using primers 5’ caaacacgtacgcgtacgatgctctagaatg and 5’ tgctatgtaacgcggaactccatatatggg and the Rac1 sequence flanked Puro-GFP cassette from pN-PITCh-GFP using primers 5’ccgcgttacatagcatcgtacgcgtacgtgtttggGGCCCAGCGAGCGGCCCTGAtgaccgagtacaagcccacg and 5’cattctagagcatcgtacgcgtacgtgtttgggACCACACACTTGATGGCCTGCAtcttgtacagctcgtccatgccgag.

Correct recombinants were verified by DNA sequencing. After transfection cells were selected using 2.5µg/ml puromycin and clones established by limiting dilution.

### Modeling

Homology models of human Sar1 and Rac1 were generated using the YASARA code (Krieger & Vriend, 2014) and the default homology modeling macro. In short, PSI-BLAST searches are made to get position specific scoring matrices from UniRef90, followed by searching the PDB for matching templates. Based on BLAST e-value and alignment scores, up to 5 templates are selected and up to 5 models generated with each template. The different models are ranked based on 1D and 3D packing. Hybrid models can then be generated by e.g. replacing loop segments and similar from the chosen main model by same parts from another model, that locally has obtained a better score.

The homology model of Sar1 is based on the Xray structure of the Sar1 dimer from *Cricetulus griseus* (PDB-ID 1F6B), containing GDP bound to the pocket. For Rac1, the main template used for the final GTP bound model is the NPH1 protein of *Avena sativa* (PDB-ID 2WKP) with N- and C-terminal segments taken from the crystal structure of human Rac1 (PDB-ID 1MH1).

The homology models of Sar1 and Rac1 were then subjected to protein-protein docking using a modified protocol of our recently developed meta-approach (Mahdizadeh *et al*, 2021). The on-line servers for the protein-protein docking engines ClusPro (Comeau *et al*, 2004; Kozakov *et al*, 2017), FireDock (Andrusier *et al*, 2007; Mashiach *et al*, 2008), Galaxy- TONGDOCK (Ko *et al*, 2012), PatchDock (Schneidman-Duhovny *et al*, 2005) and ZDOCK (Chen *et al*, 2003; Pierce *et al*, 2011) were employed using blind docking and default settings, and the 10 best scoring models from each downloaded. The 50 models were clustered based on RMSD values and evaluation of Kelley penalties (Kelley *et al*, 1996) to obtain the optimum number of clusters, using the Clustering of complexes module in the Schrödinger software [Schrödinger Release 2020-2: Maestro, Schrödinger, LLC, New York, NY, 2020.]. As consensus model was then selected the predicted complex with lowest RMSD from the centroid of the most populated cluster which in this case contained 14 out of the total predicted 50 complexes.

The obtained Rac1-Sar1 complex was analysed, and superposed onto the Sar1-Sec12 and Sar1-Sec23 crystal structures from *Saccharomyces cerevisae*, with PDB-IDs 6X90 and 1M2O, respectively. Superposition was made such that the two Sar1 units would be optimally aligned. All superposition and images were generated using the Molecular Operating Environment software [Molecular Operating Environment (MOE) 2019.01; Chemical Computing Group, Montréal, Canada, 2019].

## Acknowledgements

HF is supported by grants from the Norwegian Cancer Society (Projects 208015 & 182815), by the Research Council of Norway (Project 302452), by the Otto-Rakel-Bruun endowment and by grants from the European Commission (H2020-MSCA-ITN-860035; H2020-MSCA-ITN-859962). Funding from the Vinnova Seal-of-Excellence program 2019-02205 (CaTheDRA) to SJM is gratefully acknowledged. LAE is supported by the Swedish Research Council through grant 2019-3684. Work in the Teis lab is funded by DOC82 doc.fund, and P 29583. JCK is a recipient of a DOC Fellowship of the Austrian Academy of Sciences. The Swedish National Infrastructure for Computing is acknowledged for allocations of computing time at supercomputing center NSC, in part funded by the Swedish Research Council through grant agreement no. 2018-05973 (LAE). MGK is supported by National Institutes of Health grant ES026023. P.R.-C. Acknowledges funding from Spanish Ministry of Science and Innovation (PID2019-110298GB-I00), the European Commission (H2020-FETPROACT-01- 2016-731957), The prize “ICREA Academia” for excellence in research, Fundaci6 la Marató de TV3, and Obra Social “La Caixa”. IBEC is a recipient of a Severo Ochoa Award of Excellence from MINCIN.

## Author contributions

Conceptualization: SP, HF

Methodology: SP, HF, ALR, PRC, MJB, SG, DF

Data acquisition: SP, ED, SJM, MJB, LAE, SG, DF, VR, JCK

Data analysis, interpretation and visualization: SP, ED, HF, SJM, MJB, LAE, SG, JCK, DT

Writing - original draft: SP, HF

Writing - review & editing: SP, ED, ALR, MJB, MGK, PRC, SG, LAE, HF

## Methods

### RESOURCE AVAILABILITY

- Lead contact: For information and requests for resources please contact Hesso Farhan:
- Material availability: the generated cell line with endogenously tagged GFP-Rac1 will be made freely available.
- Data and code availability: Any information required to reanalyze the data reported in this work is available from the lead contact upon request. This work does not report any original code.

### METHOD DETAILS

#### Cell culture and treatment

HeLa cells were maintained in DMEM (Lonza, #12-604F) supplemented with 10% FCS (Life Technologies, #10500064), 50 units/ml penicillin and 50 μg/ml streptomycin (Lonza, #DE17- 603E), in a humidified 5% CO2 atmosphere at 37 °C. PC3 cells and MDA-MB-231 cells were cultured in RPMI medium with supplements as for HeLa cells. CRISPR Rac1 knockout PC3 cells were extensively characterized previously (Baker et al., 2020).

For all Rac1 inhibition experiments, cells were treated with 50 μM NSC23766 or 0.1% (v/v) DMSO in complete medium for 4 h. For actin disruption experiments, cells were treated with 0.5, 1, 1.5 and 2 µM cytochalasin D or latrunculin A in complete medium for 15 and 30 min. For RUSH assays, cells were preincubated with 1 µM cytochalasin D or latrunculin A for 15 min before starting the cargo release with biotin.

siRNA oligos were delivered to the cells using HiPerFect transfection reagent. Briefly, 10 – 20 nM siRNA duplexes were complexed with the transfection reagent in 100 µl serum and antibiotics free cell culture medium, incubated at room temperature for 5 min and added to the cells immediately after seeding for experiments. Knockdown efficiency after 72 h was measured to check that the levels of desired proteins were reduced. For Rac1 knockdown experiments, the siGenome non-targeting siRNA pool from Dharmacon were used as control. The AllStars negative control siRNA from Qiagen was used as control for INF2 knockdown experiments.

#### FRAP microscopy

FRAP analysis was carried out as described previously(Centonze et al., 2019; Farhan et al., 2010).

#### Preparation of polydimethylsiloxane (PDMS) membranes

Stretchable PDMS (Sylgard Silicone Elastomer Kit, Dow Corning) membranes were prepared as previously described(Kosmalska et al., 2015). Briefly, a mix of 10:1 base to crosslinker ratio was spun on plastic wafers for 1 minute at 500 rpm and then cured overnight at 65 °C. Once polymerized, membranes were peeled off the wafers and assembled onto a metal ring for fibronectin coating, cell seeding and microscopy experiments.

#### Mechanical stimulation of the cells and live cell imaging

Cell mechanical stimulation was done as previously described(Kosmalska *et al*., 2015). Briefly, a 150 μL droplet of a 10 μg/mL fibronectin solution (Sigma, #F0895) was deposited in the center of the membrane mounted in the ring. After overnight incubation at 4 °C, the fibronectin solution was rinsed. HeLa cells expressing GFP-Sec16A were then seeded on the fibronectin coated membranes and allowed to attach for up to 1 h. Then ring-containing membranes were mounted in the stretch system previously described(Kosmalska *et al*., 2015). Live-cells were subjected to a 7.5 % equibiaxial strain. Images of the cells were acquired with 500 ms interval frames, with a 60x objective (NIR Apo 60X/WD 2.8, Nikon) mounted in an inverted microscope (Nikon Eclipse Ti) with a spinning disk confocal unit (CSU-W1, Yokogawa), a Zyla sCMOS camera (Andor) and using the Micromanager software.

#### Live cell imaging and optogenetics

Live cell imaging of HeLa cells co-expressing mCherry-Rac1 and GFP-Sec61 or mCherry-Rac1 and GFP-Sec16 was performed on Andor Dragonfly spinning disk equipped with a Nikon Ti2E inverted optical microscope (60 × TIRF objective (Plan-APOCHROMAT 60 × /1.49 Oil) and an EMCCD camera (iXon Ultra 888, Andor). Cells were maintained at 37 °C in 5% CO2 in a heating chamber (Oko lab). SRRF-Stream mode in Fusion (version 2.1, Andor) was used for image acquisition with an additional 1.5 × magnification. Following imaging parameters were used. SRRF Frame count: 150-200, SRRF Radiality Magnification: 4×, SRRF Ring Radius: 1.4 px, SRRF Temporal Analsysis: Mean and SRRF FPN correction: 75 -100 frames.

For optogenetics, HeLa cells expressing CIBN-GFP-Sec61β, Cry2-TIAM-mCherry and SNAP- Sec16A were stimulated for 30 sec with a 488 nm laser.

#### Retention using selective hooks (RUSH) assay

A step-by-step protocol for RUSH assay has been described elsewhere(Boncompain and Perez, 2012). We used HeLa cells stably expressing the Golgi resident protein Mannosidase II in RUSH system, and monitored its arrival at the Golgi for up to 30 minutes in our experiments.

#### ERES, RUSH quantification and colocalization analysis

ERES and RUSH quantification, and colocalization analysis were all carried out using Fiji(Schindelin et al., 2012). Prior to analysis, background was subtracted from the immunofluorescence images using “subtract background” command in Fiji. To obtain ERES count in cells, images were thresholded (Otsu algorithm) and ERES were counted using “Analyze particle” command. Counts were then normalized to total cells.

For quantification of RUSH experiments, ManII intensity in the Golgi was measured by drawing a ROI. To measure extra Golgi intensity of ManII, the ROI was then transferred outside the Golgi. Ratio of ManII at the Golgi to extra Golgi region was calculated for data visualization. The fraction overlap of Rac1-Sar1 complex with Sec31 was calculated using DiANA(Gilles et al., 2017) plugin in Fiji.

#### Cell lysis, immunoprecipitation and immunoblotting

For immunoprecipitation experiments, cells were lysed in immunoprecipitation buffer (20 mM Tris-HCl, pH 7.4, 150 mM NaCl, 1 mM MgCl2, 10% glycerol, 0.5% NP-40) and GFP-trap (ChromoTek, #gta-100) was performed following manufacturer’s protocol.

#### Active Sar1 pulldown assay

To the microsomal fractions, 750 ng of recombinant Rac1 or Sar1 or both and GTP or GTPγS (100 µM final concentration) were added. The reaction mixture was then incubated at 30 °C with constant agitation for 30 min for GTP-loading prior to active Sar1 pulldown with anti-active Sar1 monoclonal antibody provided in the kit.

#### Rac1 GFP tagging

Rac1 was gene edited using a modified PITCh technology (Sakuma et al., 2016) adjusted for generating N-terminal fusion proteins (Lin et al., 2019). A Rac1 specific gRNA 5’ ACACTTGATGGCCTGCATCA was cloned into plasmid pX330-BbsI-PITCh (Addgene plasmid #127875) and transfected along with pN-PITCh-GFP-Rac1 into HeLa cells using JetPrime (Polyplus, Illkirch, France). Plasmid pN-PITCh-GFP-Rac1 contains a Puromycin resistance-T2A- eGFP cassette flanked by 50 bp of genomic RAC1 sequence flanking the CRSIPR-CAS9 induced DNA double strand break. This plasmid was constructed by HiFi- mediated in vitro recombination (NEB) of two PCR based fragments generated by amplifying the pN-PITCh-GFP (Addgene plasmid #127888) vector backbone using primers 5’ caaacacgtacgcgtacgatgctctagaatg and 5’ tgctatgtaacgcggaactccatatatggg and the Rac1 sequence flanked Puro-GFP cassette from pN- PITCh-GFP using primers 5’ ccgcgttacatagcatcgtacgcgtacgtgtttggGGCCCAGCGAGCGGCCCTGAtgaccgagtacaagcccacg and 5’ cattctagagcatcgtacgcgtacgtgtttgggACCACACACTTGATGGCCTGCAtcttgtacagctcgtccatgccgag. Correct recombinants were verified by DNA sequencing. After transfection cells were selected using 2.5µg/ml puromycin and clones established by limiting dilution.

#### Modeling

Homology models of human Sar1 and Rac1 were generated using the YASARA code (Krieger and Vriend, 2014) and the default homology modeling macro. In short, PSI-BLAST searches are made to get position specific scoring matrices from UniRef90, followed by searching the PDB for matching templates. Based on BLAST e-value and alignment scores, up to 5 templates are selected and up to 5 models generated with each template. The different models are ranked based on 1D and 3D packing. Hybrid models can then be generated by e.g. replacing loop segments and similar from the chosen main model by same parts from another model, that locally has obtained a better score.

The homology models of Sar1 and Rac1 were then subjected to protein-protein docking using a modified protocol of our recently developed meta-approach (Mahdizadeh et al., 2021). The on- line servers for the protein-protein docking engines ClusPro (Comeau et al., 2004; Kozakov et al., 2017), FireDock (Andrusier et al., 2007; Mashiach et al., 2008), Galaxy-TONGDOCK (Ko et al., 2012), PatchDock (Schneidman-Duhovny et al., 2005) and ZDOCK (Chen et al., 2003; Pierce et al., 2011) were employed using blind docking and default settings, and the 10 best scoring models from each downloaded. The 50 models were clustered based on RMSD values and evaluation of Kelley penalties (Kelley et al., 1996) to obtain the optimum number of clusters, using the Clustering of complexes module in the Schrödinger software [Schrödinger Release 2020-2: Maestro, Schrödinger, LLC, New York, NY, 2020.]. As consensus model was then selected the predicted complex with lowest RMSD from the centroid of the most populated cluster which in this case contained 14 out of the total predicted 50 complexes.

The obtained Rac1-Sar1 complex was analysed, and superposed onto the Sar1-Sec12 and Sar1- Sec23 crystal structures from *Saccharomyces cerevisae*, with PDB-IDs 6X90 and 1M2O, respectively. Superposition was made such that the two Sar1 units would be optimally aligned. All superposition and images were generated using the Molecular Operating Environment software [] Molecular Operating Environment (MOE) 2019.01; Chemical Computing Group, Montréal, Canada, 2019].

## Supplementary Figures

**Fig. S1.**
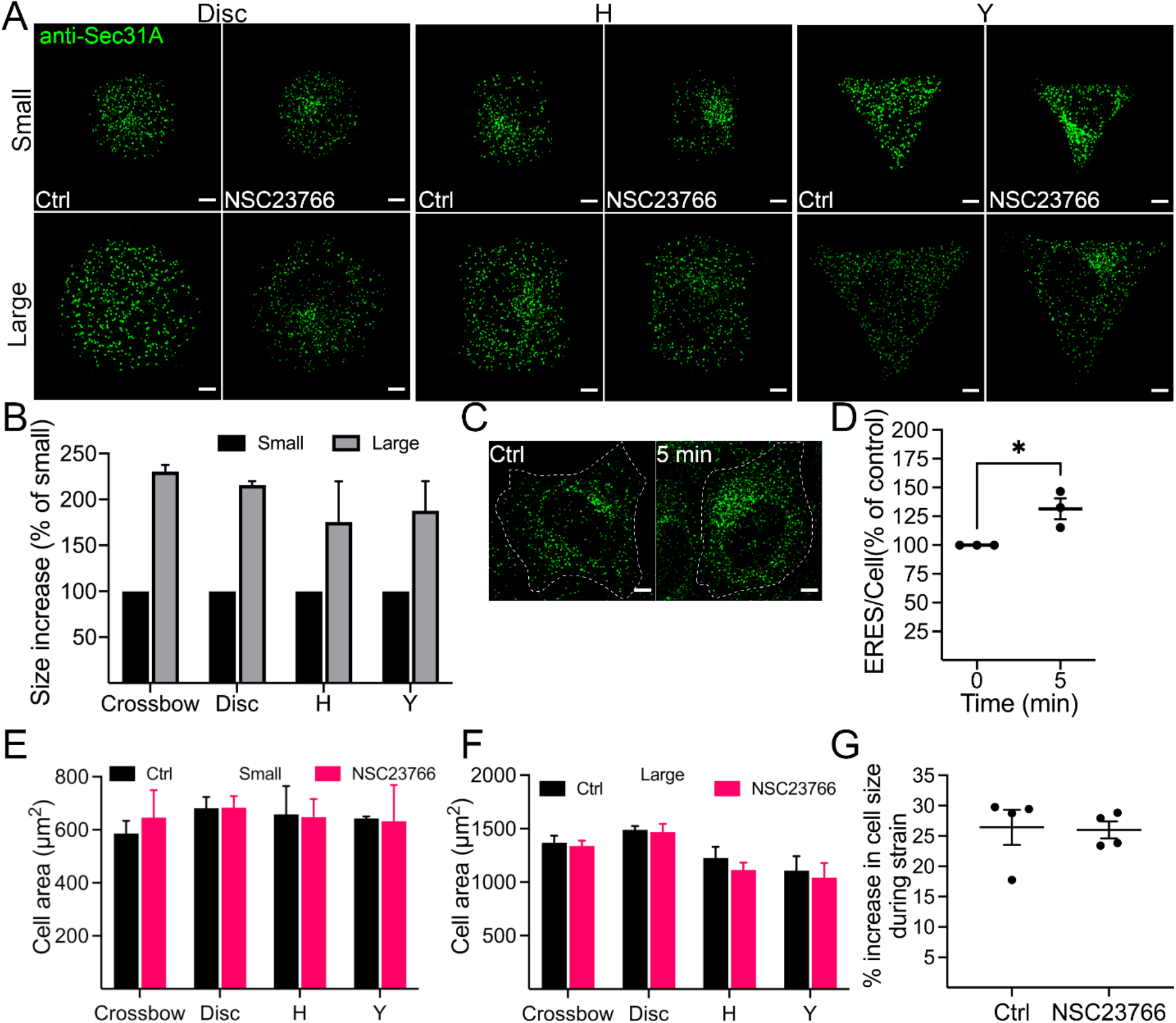
(related to Figure 1) (A) Representative images showing Sec31A labelled ERES in HeLa cells seeded on fibronectin coated micropatterned chips of different geometries (Disc, H and Y) and different sizes (small or large). Cells were treated with either DMSO (Ctrl) or Rac1 inhibitor (NSC23766) for 4 h prior to immunolabelling with Sec31A. Scale bars: 5 µm. (B) Quantification graph showing increase in area covered by HeLa cells that were covering the large geometries (Crossbow, Disc, H or Y) on fibronectin coated micropatterned chips. For better visualization, data are expressed as percentage of area covered by cells on the small geometries. Error bars represent standard deviation. 50-60 cells from two independent experiments were counted. (**C** and **D**) HeLa cells were subjected to osmotic shock for the indicated time point and changes in ERES was monitored by counting Sec31A marked ERES. (C) Representative immunofluorescence images showing ERES in control cells and cells subjected to osmotic shock. (D) Quantification of ERES in C. At least 30 cells were counted in each experiment (n = 3). Mean ± SEM and dots representing average ERES/cell values from each experiment are shown. * = P- value < 0.05 (**E** and **F**) Graphs show area covered by HeLa cells treated with DMSO (Ctrl) or Rac1 inhibitor (NSC23766) grown on small (E) or large (F) fibronectin coated micropatterned chips. Between 50-60 cells were counted in two experiments. Values show average cell area ± standard deviation. (**G**) Quantification graph showing increase in size of HeLa cells during 7.5% biaxial strain. A total of 19 cells were quantified in three live cell imaging experiments. Cell size before strain was used to calculate percent increase in cell size during strain. Mean ± SEM and dots representing average values from each experiment are shown.

**Figure S2.**
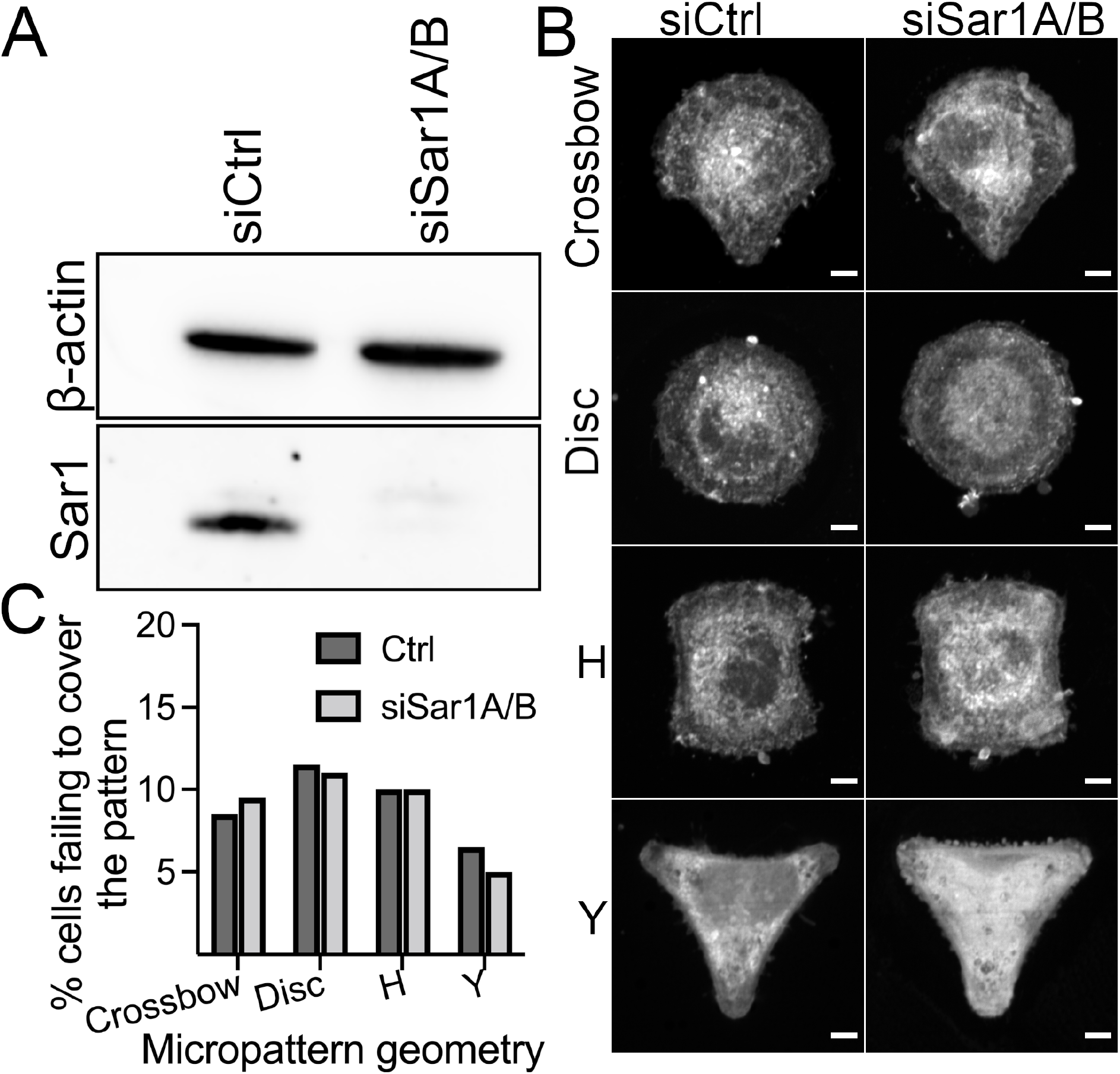
(A) HeLa cells were co-transfected with 5 nM siSar1A and 5 nM siSar1B (siSar1A/B) for 48 h. Control cells (siCtrl) were transfected with 10 nM non-targeting siRNA oligos. The efficiency of knockdown was evaluated by Western blot. A representative immunoblot showing the knockdown efficiency of siRNA oligos. (B) Representative immunofluorescence images of HeLa cells cultured on micropatterned surface of multiple geometries of small size (600 µm^2^). Cells were treated as described in (A), and after 48 h, cells were trypsinized and 80,000 cells were seeded on micropatterned chips. Cells were allowed to attach for 3 h, stained with CellMask for 5 min and processed for microscopy. Scale bars: 5 µm (**B**) Quantification graph showing the percentage of cells failing to entirely cover the different geometries (described in B). Between 60-70 cells were imaged in two independent experiments.

**Fig. S3.**
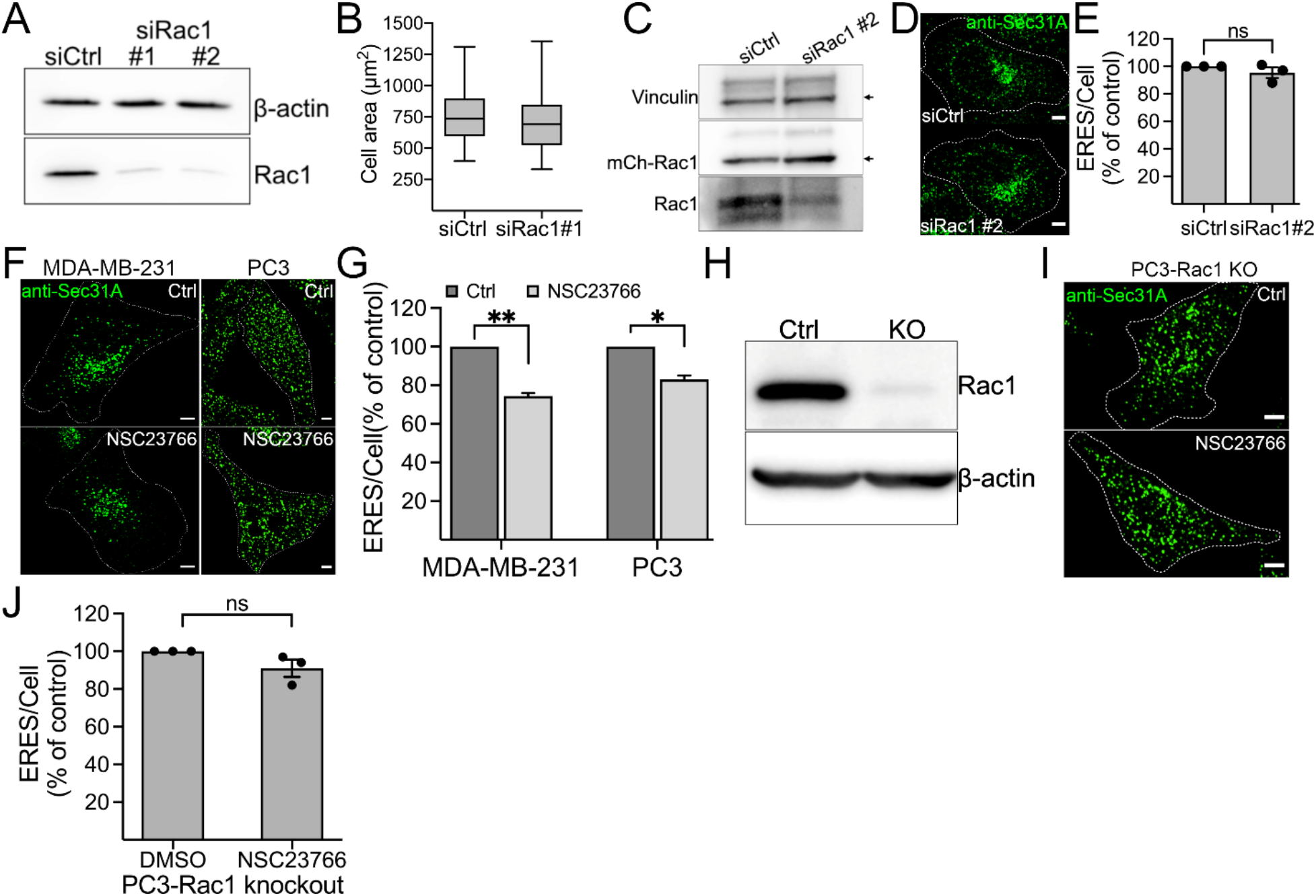
(A) HeLa cells were transfected with two siRNAs against Rac1 (siRac1 #1 and siRac1 #2) for 72 h. The efficiency of knockdown was evaluated by Western blot. A representative immunoblot showing the knockdown efficiency of siRNA oligos. (B) Area covered by HeLa cells transfected with non-targeting control siRNA (siCtrl) or siRNA against Rac1 (siRac1 #1). Number of cells = 30, number of experiments = 3 (**C-E**) HeLa cells stably expressing mCherry-tagged siRNA resistant Rac1 was transfected with control siRNA (siCtrl) or siRNA against Rac1 (siRac1 #2) for 72 h. Cells were then lysed for immunoblotting or processed for immunostaining with Sec31A. (**C**) A representative immunoblot showing reduction in the levels of endogenous Rac1 while siRNA resistant mCherry-Rac1 remains unaffected. (**D**) Immunofluorescence images showing ERES in siCtrl and siRac1 treated cells. Scale bars, 5 µm. (**E**) Quantitation of experiment in C. Graph shows mean ± SEM (n = 3, siCtrl: 117 cells, siRac1 #2: 94 cells). ns = not siginificant (**F** and **G**) Inhibition of Rac1 with NSC23766 (50 µM for 4 h) reduces ERES in MDA-MB-231 and PC3 cells. Sec31A marked ERES are shown in E and quantification of ERES is shown in F. Scale bars: 5 µm. (**G**) Graph shows mean ± SEM (n = 3 experiments, at least 30 cell/experiment), where asterisks (*) represent statistical significance at p-value < 0.05 (Student’s unpaired *t* test). (**H**) Immunoblot of CRISPR Rac1 knockout PC3 cells. (**I** and **J**) Effect of Rac1 inhibitor NSC23766 is on-target. CRISPR Rac1 knockout PC3 (PC3-Rac1 KO) cells were treated with DMSO (Ctrl) or NSC23766 (50 µM) for 4 h and ERES was quantified by counting Sec31A labelled puncta from confocal images. (H) Representative images showing ERES in PC3-Rac1 KO Ctrl or NSC23766 treated cells. Scale bars, 5 µm. (I) Treating PC3-Rac1 KO cells with NSC23766 does not reduce ERES. Average ERES per cell are displayed in % of control (n = 3, number of cells = 60-90). ns = not significant

**Fig. S4.**
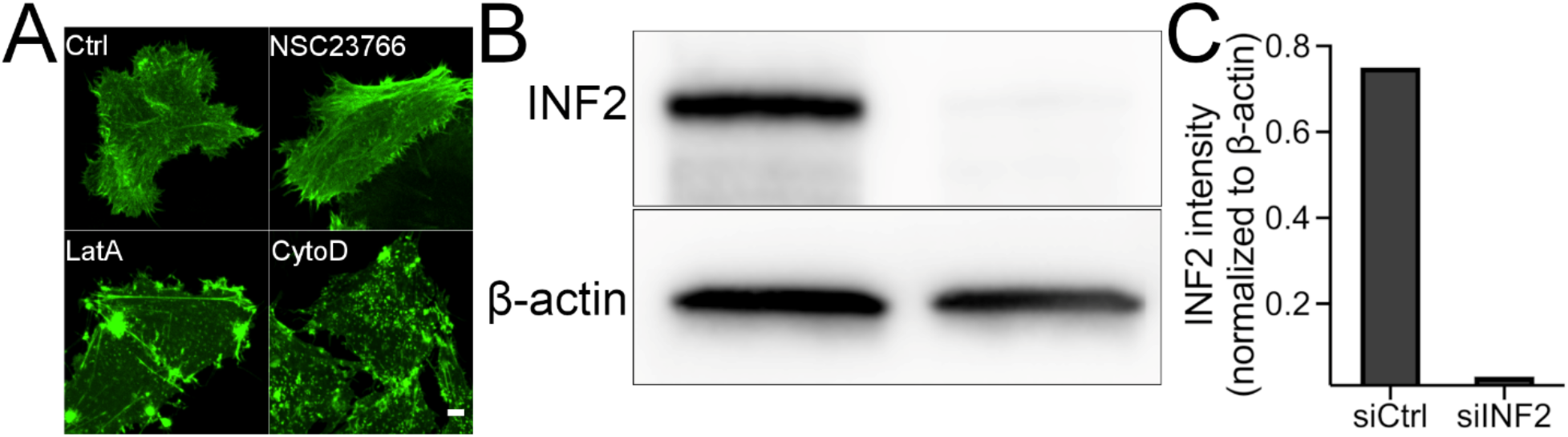
(**A**) Treatment of HeLa cells expressing GFP-tagged actin chromobody with 0.5 µM cytochalasin D (CytoD) or latrunculin A (LatA) for 15 min severely affects actin compared to DMSO (Ctrl) or Rac1 inhibitor (NSC23766, 50 µM for 4 h). (**B**-**C**) Immunoblot (B) and quantification graph (C) showing reduction in endogenous INF2 level in HeLa cells following transfection with 25 nM siRNA targeting INF2 (siINF2) for 72 h.

**Fig. S5.**
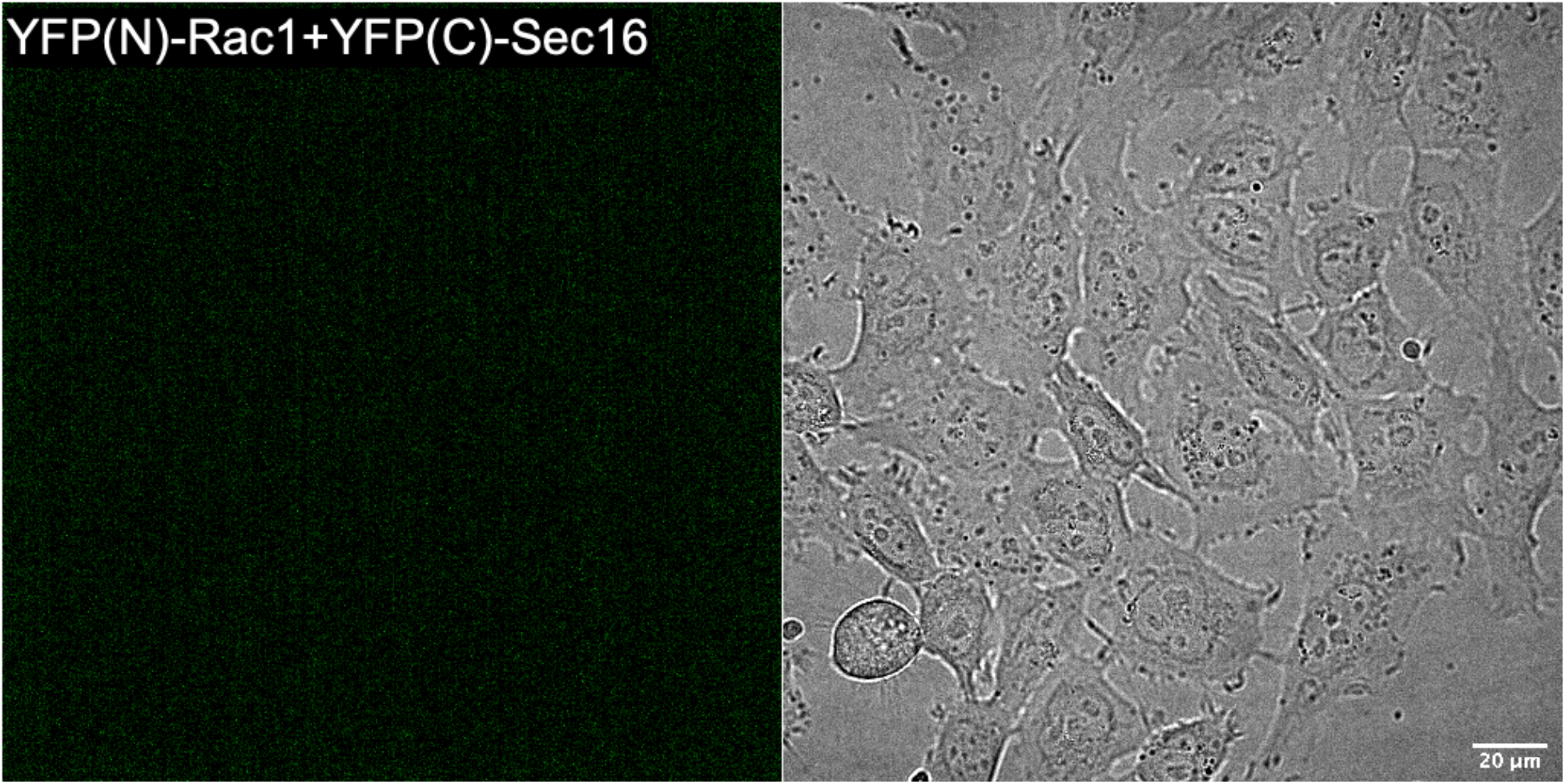
YFP(N)-Rac1 and YFP(C)-Sec16 co-transfected HeLa cells. Lack of positive YFP signal suggest no interaction of Rac1 and Sec16.

**Movie S1.**

ERES dynamics in HeLa cells before and during equibiaxial strain. Timelapse movie corresponding to Fig. 1C. Frames were captured every 500 ms.

**Movie S2.**

ManII-RUSH trafficking in HeLa cells seeded on PDMS membranes. Timelapse movie corresponding to Fig. 1H. Frames were captured every 90 sec over the course of 30 min.

**Movie S3.**

A transient pool of Rac1 at the ER. Timelapse movie corresponding to Fig. 3A, showing co- occurence of Rac1 (magenta) with ER (labelled with GFP-Sec61β). 15 sec/frame for 3 min 45 secs.

**Movie S4.**

Movie corresponding to Fig. 5A shows Rac1 (magenta) co-occuring with GFP-Sec16 labelled ERES. Frames were acquired every 5 sec over the course of 75 secs.

## KEY RESOURCES TABLE

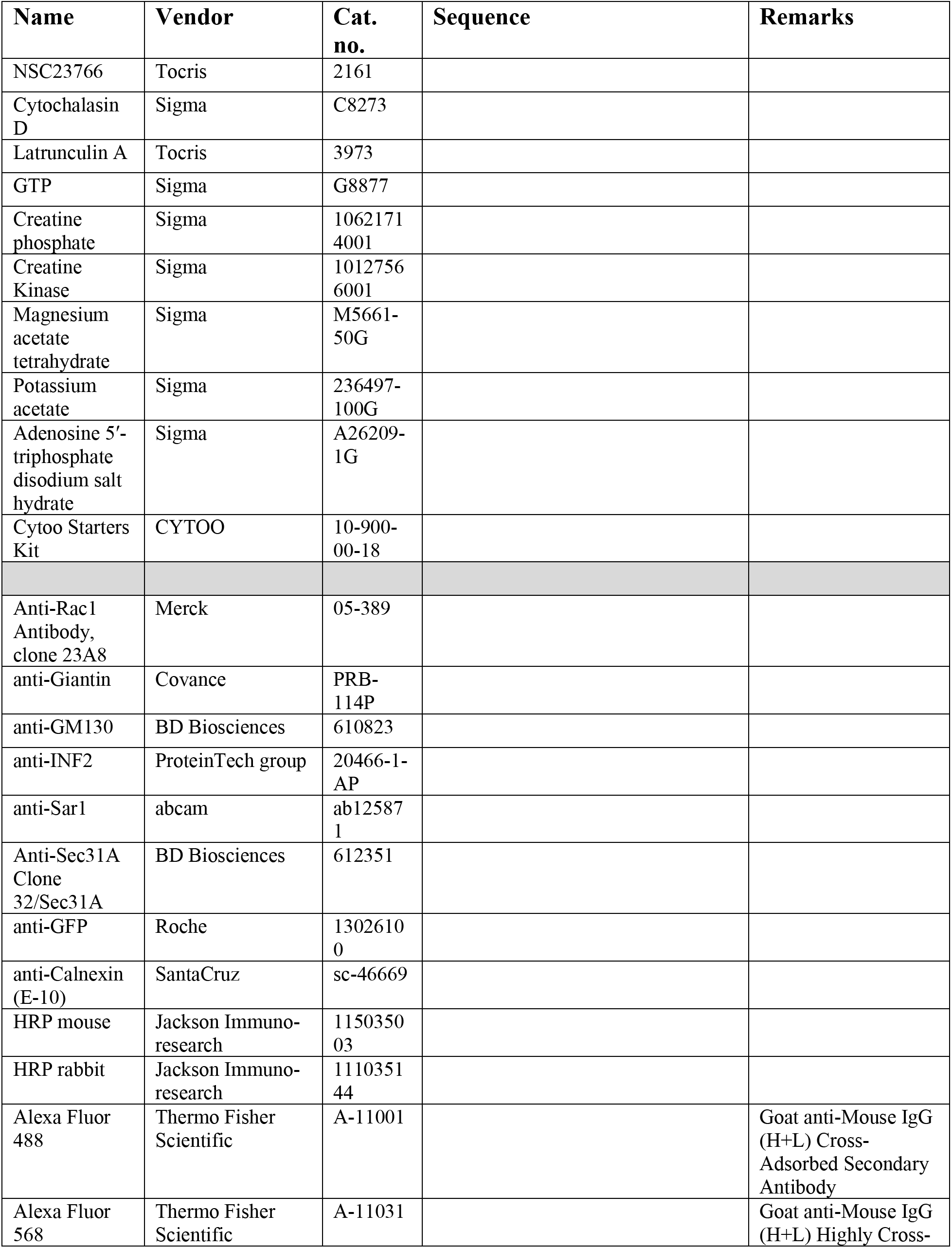
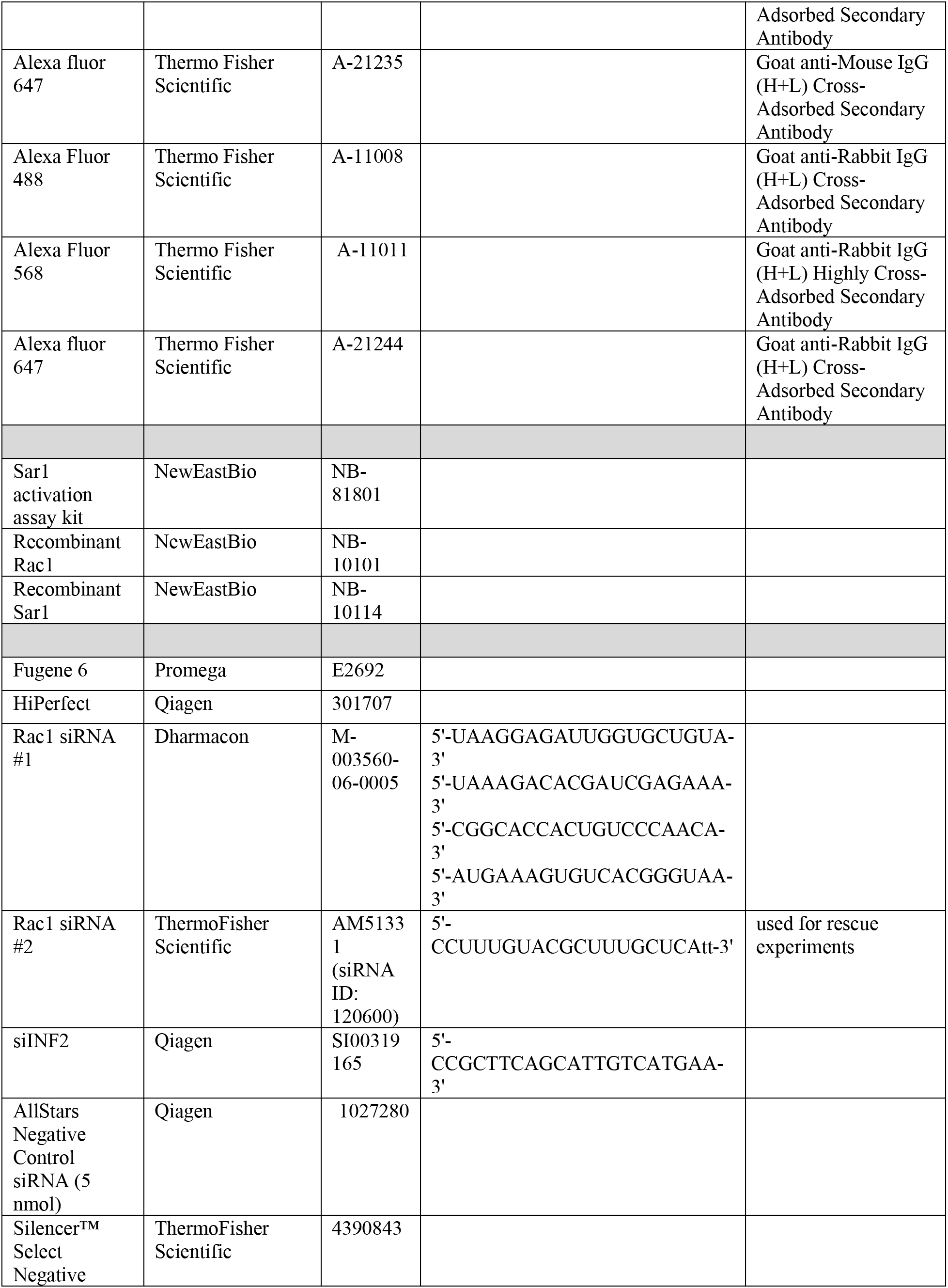
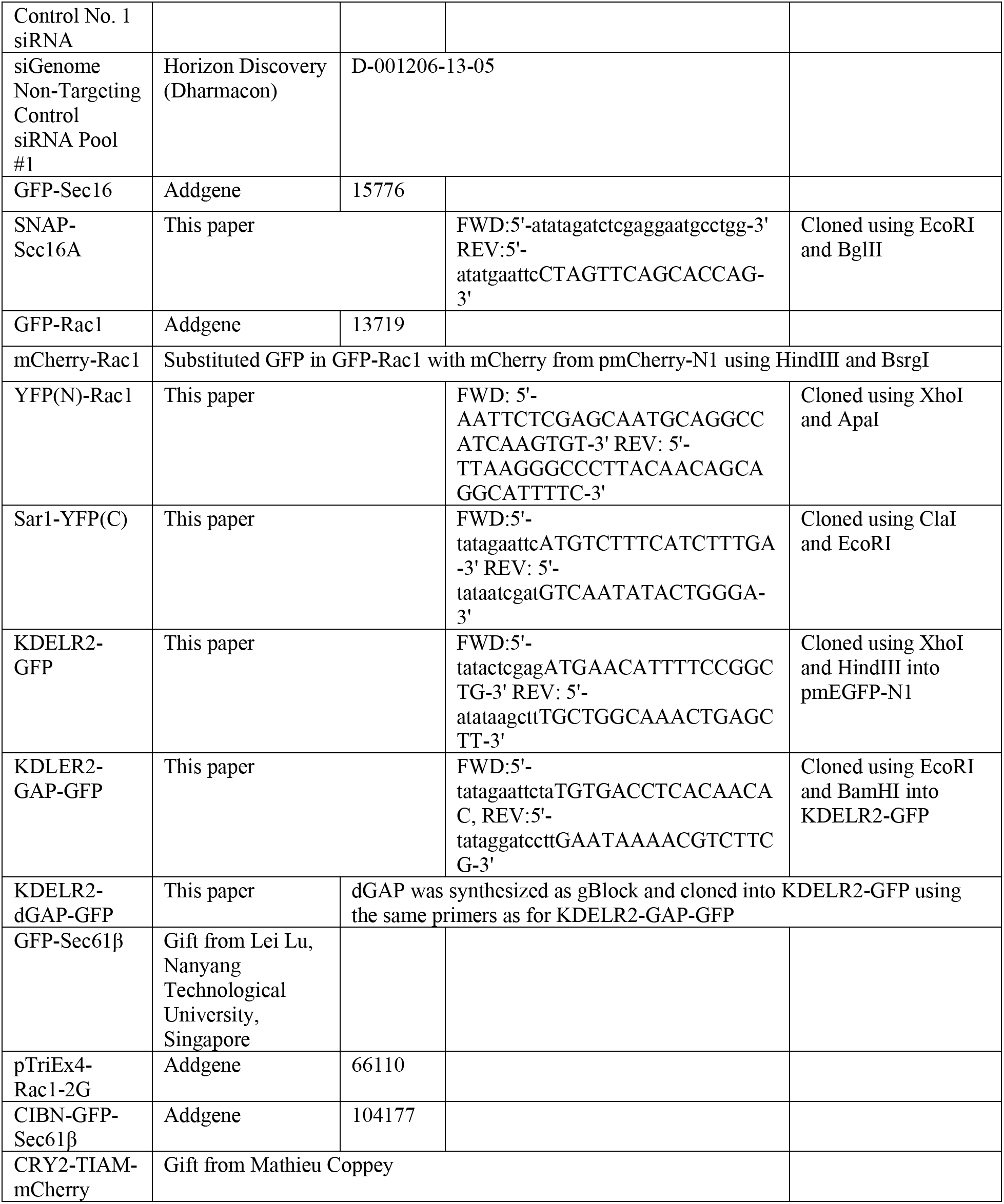

